# Tracing the genetic diversity of the bread wheat D genome

**DOI:** 10.1101/2024.08.23.609385

**Authors:** Zihao Wang, Wenxi Wang, Yachao He, Xiaoming Xie, Zhengzhao Yang, Xiaoyu Zhang, Jianxia Niu, Huiru Peng, Yingyin Yao, Chaojie Xie, Mingming Xin, Zhaorong Hu, Qixin Sun, Zhongfu Ni, Weilong Guo

## Abstract

Bread wheat (*Triticum aestivum*) became a globally dominant crop after incorporating the D genome from donor species *Aegilops tauschii*, while evolutionary history shaping the D genome during this process remains elusive. Here, we proposed a renewed evolutionary model linking *Ae. tauschii* and hexaploid wheat D genome, based on an ancestral haplotype map covering a total of 762 *Ae. tauschii* and hexaploid wheat accessions. We dissected the evolutionary process of *Ae. tauschii* lineages and clarified L3 as the most ancient lineage. A few independent intermediate accessions were reported, demonstrating the low-frequent inter-sublineage geneflow enriched the diversity of *Ae. tauschii*. We discovered that the D genome of hexaploid wheat inherited from a unified ancestral template, but with a mosaic composition that is highly mixed by three *Ae. tauschii* L2 sublineages located in the Caspian coastal region, suggesting the early agricultural activities facilitate the innovation of D genome compositions that finalized the success of hexaploidization. We further found that the majority (65.6%) of polymorphisms were attributed to novel mutations absent during the spreading of bread wheat, and also identified large *Ae. tauschii* introgressions from wild *Aegilops* lineages, expanding the diversity of wheat D genome and introducing beneficial alleles. This work decoded the mystery of the wheat hexaploidization process and the evolutionary significance of the multi-layered origins of the genetic diversity of the bread wheat D genome.

## Introduction

Maximizing the genetic potential of crops needs a comprehensive understanding of the genetic diversity in extant germplasm, while the formation of which is deeply shaped by the evolution and domestication history^1,2^. As a hexaploid crop formed only 8,500-9,000 years ago, bread wheat (*Triticum aestivum*) expanded its habitat from a small core area within the Fertile Crescent to diverse environments worldwide and played a pivotal role in the global food system^3^. This wide adaptability is partially ensured by its genome plasticity after gaining the D subgenome ^4^. The addition of the D subgenome also provides bread wheat with superior quality and grain yield than tetraploid wheat. Nonetheless, the evolutionary process leading to the bread wheat D genome is not fully understood^5^, which hindered the efficiency of giving possible solutions for future challenges in food security.

An in-depth understanding of the origin of the hexaploid wheat D genome relies on its evolutionary relationship with wild *Aegilops tauschii*, the D genome donor, while the population structure of *Ae. tauschii* remains a discrepancy. It could be partially explained by the important role of cross-pollination in the evolution of wild goatgrass, despite being mostly self-pollinating species^6-9^. *Ae. tauschii* had at least two major subspecies or two lineages, *tauschii* and *strangulata* (hereafter L1 and L2)^10,11^, naturally spreading from southeastern Turkey to western China with its center of genetic diversity in the southern Caspian coastal region^12^, while the existence of the third putative lineage (hereafter L3) remained controversial^13-15^. Furthermore, the evolutionary relationship among genetically distinct sublineages of *Ae. tauschii* is also vital in characterizing the origin of the hexaploid wheat D genome from *Ae. tauschii*.

The bread wheat D genome diversity is only approximately 16% of that of the A and B genomes^16,17^. This lower level of variation has seriously restricted the selection of desirable agronomic traits in wheat breeding^18,19^. It has been proposed that this limitation of genetic diversity resulted from a polyploidy bottleneck and that only a small population of *A. tauschii* contributed to the formation of bread wheat^20^. On the contrary, highly diversified genomic regions existed between the hexaploid wheat D genome and each of the available *Ae. tauschii* genomes^13,21^. The extant evolutionary model was unable to settle on a consensus on the origin of the hexaploid wheat D genome. Furthermore, genetic and genomic studies have reported the differential presence of L2 and L3 segments in the hexaploid wheat D genome^22-25^. A genomic study showed that *Ae. tauschii* L3 also contributed ∼ 1% on average to the hexaploid wheat D genome, based on which the multiple hybridization origin hypothesis of bread wheat has been proposed^14^, though the influencing of introgression remains uncertain. Encouragingly, since entering the genomic era from wild goatgrass^26-28^ and hexaploid wheat^29,30^, sequencing data has supported to propose of more preferable insights about the evolution of D diversity^31,32^. Nevertheless, the evolutionary relationship between bread wheat D genome and its donor species *Ae. tauschii*, together with the formation process of bread wheat D genome genetic diversity, remains sketchy.

Here, we present the most comprehensive pan-ancestry *Triticum-Aegilops* D genome haploblock map and study its significance in the evolution of the D genome. We constructed the map based on 762 *Ae. tauschii* and hexaploid wheat accessions and precisely trace the origin of genetic diversity from hexaploid wheat D genome back to *Ae. tauschii*. Based on cytoplasmic and nuclear evidence, we confirmed the existence of an ancient lineage and independent intermediate accessions, which reconciled the incongruity in the subspecies delimitation of *Ae. tauschii*. Genome-wide haplotype analysis discovered a conserved ancestral genomic template across the hexaploid wheat D genome. By tracing the origin of this genomic template, we showed this template was a mosaic composition contributed by the admixture of three *Ae. tauschii*. Patterns of ancestral haplotypes revealed large blocks introgressed from *Ae. tauschii* lineages after hexaploidization, accumulated along with independent western and eastern spreading from the origin of hexaploid wheat. By excluding the inherited diversity from *Ae. tauschii*, we further demonstrated that novel mutations contributed nearly 2/3 of the diversity of the hexaploid wheat D genome. Our results unravel the complex evolutionary history of the bread wheat D genome and offer a reference for the precious utilization of D genome diversity in future genetic improvement.

## Results

### The pan-ancestry haploblock map of the *Triticum-Aegilops* D genome

By constructing the pan-ancestry haploblock map of the A&B genome of polyploid wheat, handling the complex architecture and frequent variations, we revealed the dispersed emergence and protracted domestication of polyploid wheat, which proved the haploblock map helpful for dissecting the evolution of wheat^33^. However, the haploblock map of the D genome still lacks. To dissect the origin and evolution of the wheat D genome by constructing the pan-ancestry haploblock map, we collected a panel of whole-genome sequencing data of accessions from genera *Triticum* and *Aegilops*, including 470 *Aegilops tauschii* and 292 hexaploid wheat accessions, representing its natural geographic distribution **(Supplementary Table 1**)^13,14,34,35^. Based on ∼76.9 million single nucleotide polymorphisms (SNPs), we identified ancestral haplotype groups (AHGs) and constructed the pan-ancestry haploblock map of the *Triticum-Aegilops* D genome **(Supplementary Fig. 1)**. The saturation curve estimated from D genome-wide AHG types confirmed that the majority of genetic diversity in *Ae. tauschii* has been captured in the map **(Supplementary Fig. 2)**. There is a median of 22 AHG types in *Ae. tauschii* across D chromosomes, while AHG has been fixed as only one dominant type in hexaploid wheat among most genomic regions, and 84.9% of the hexaploid wheat AHGs could be traced back to collected *Ae. tauschii* accessions.

**Fig. 1.**
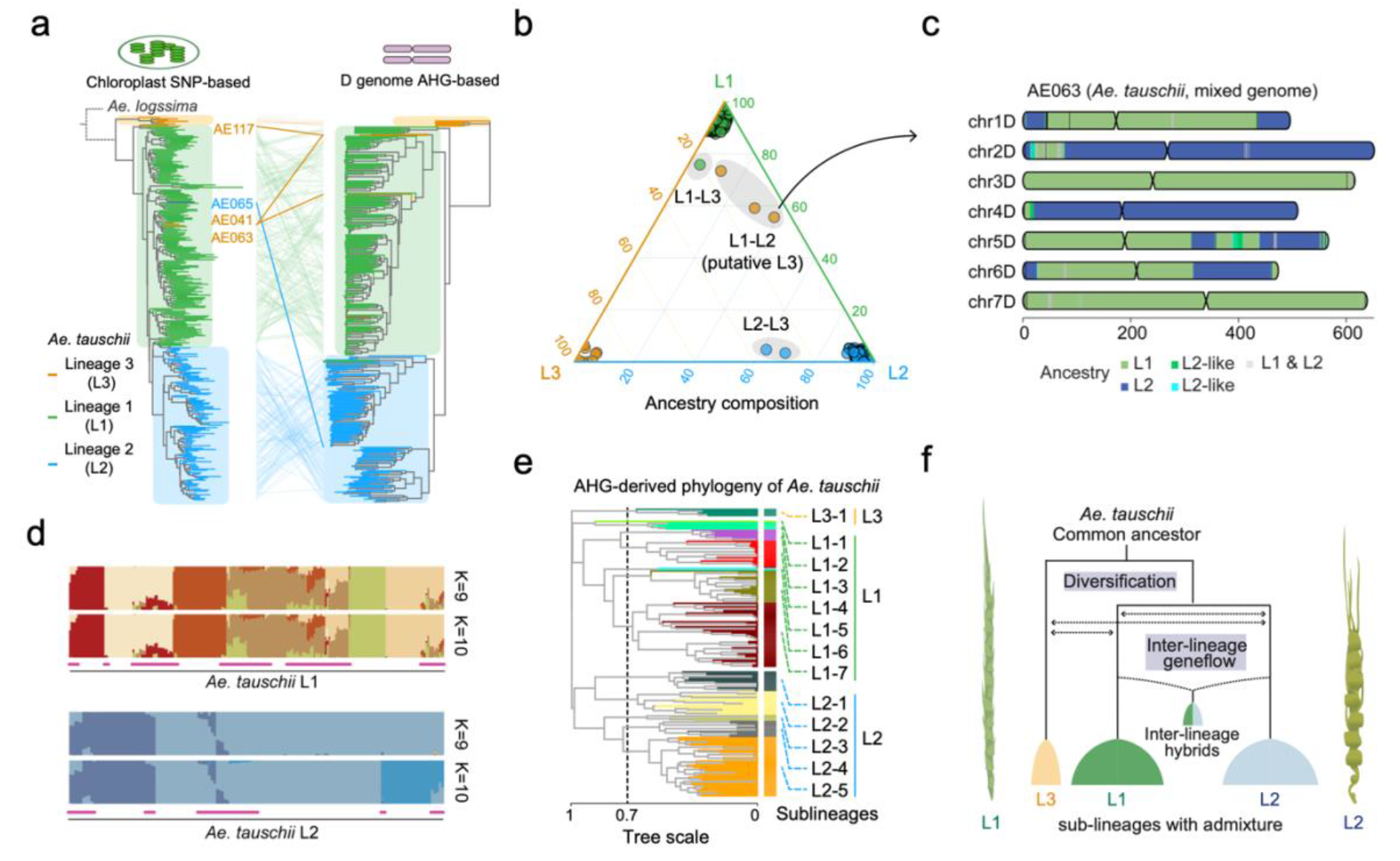
Phylogenetic relationship between *Ae. tauschii* sublineages. **a**, Comparison between the chloroplast SNP-based neighbor-joining tree (left) and D genome AHG-derived distance neighbour-joining tree (right) of *Ae. tauschii*. The chloroplast tree is rooted by assigning *Ae. speltoides* accessions as the outgroup. The same accession in cytoplasmic tree and genomic tree is connected with colored lines. Labelled accessions indicate they were clustered to a clade different from its delimitated lineages. **b**, Ternary plot shows relative L1, L2 and L3 ancestral ratios of all *Ae. tauschii* accessions. Six accessions show high ratios of inter-lineage ancestry (noted with shaded circle), including three previously putative *Ae. tauschii* L3 accessions. **c**, Chromosomal composition of ancestral haplotypes for previously putative *Ae. tauschii* L3 accession AE063, which is of the intermediate lineage of *Ae. tauschii* L1 and L2. **d**, Individual ancestry coefficients of *Ae. tauschii* accessions in L1 and L2 lineages inferred using ADMIXTURE (K = 9, 10). Purple lines, potential intermediate accessions (ancestral coefficients < 99%). **e**, Sublineage delimitation of non-admixture *Ae. tauschii* accessions based on neighbor-joining tree constructed with D genome AHG-derived distance. Colors distinguish 13 *Ae. tauschii* sublineages. **f**, Refined evolutionary model of three *Ae. tauschii* sublineages highlight L3 as the acient lineage and inter-lineages gene flows.

**Fig. 2.**
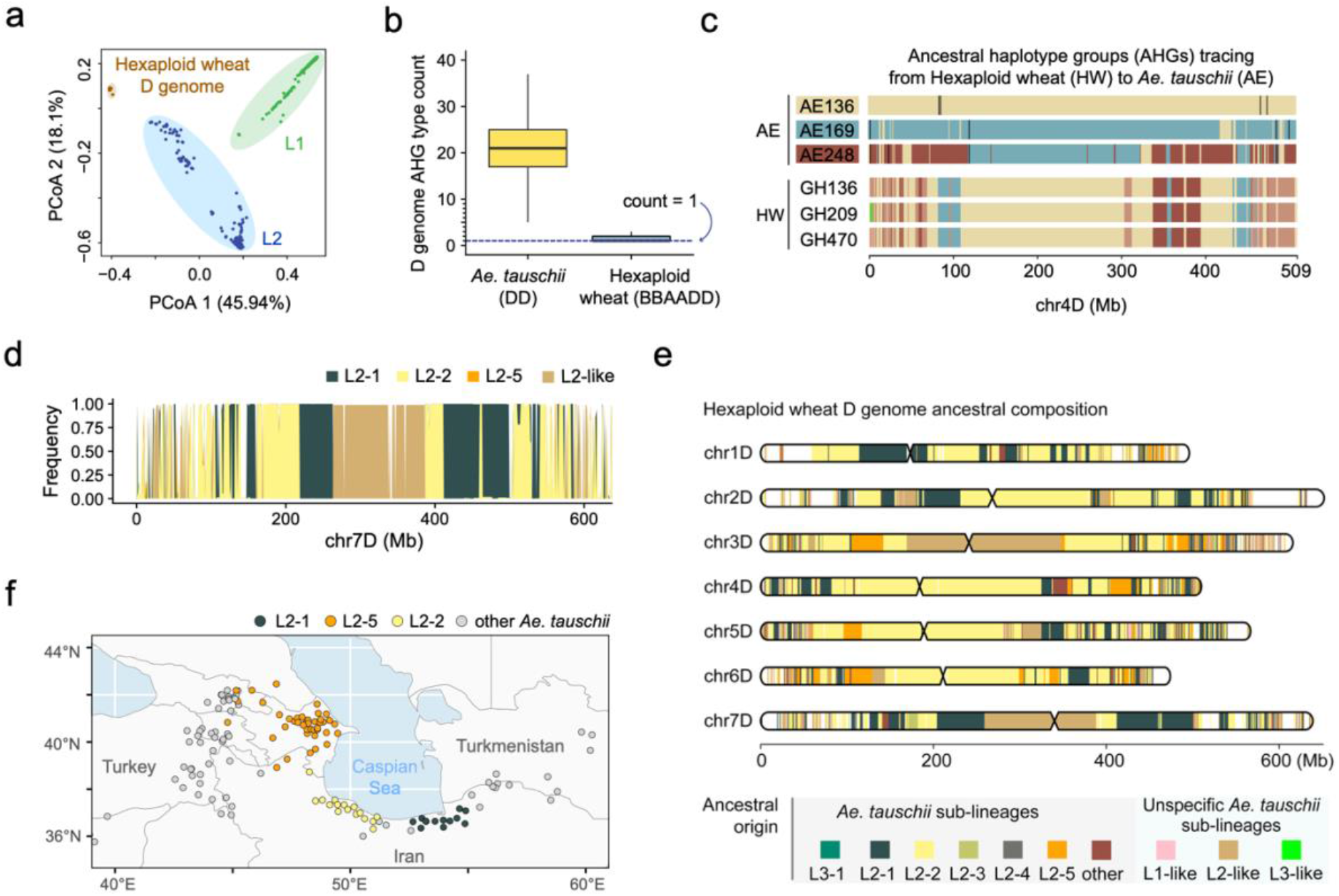
Origin of hexaploid wheat D genome ancestral temple from *Ae. tauschii*. **a**, Principal component analysis (PCoA) of *Ae. tauschii* L1, L2 and hexaploid wheat accessions, based on AHG-derived distances in D genomes. Window size, 1-Mb. **b**, Comparison of the of AHG type polymorphism in D genome between *Ae. tauschii* and hexaploid wheat. **c**, Colored mosaic graphs of AHGs across chromosome 4D for three representative *Ae. tauschii* (AE) accessions and three hexaploid wheat (HW) accessions. Priority order, AE > HW, and AE136>AE169>AE248. Colors distinguish AE accession-dominated AHGs. **d**, The AHG frequency distribution of hexaploid wheat across 7D chromosome, by tracing back to sublineages of *Ae. tauschii* L2, include L2-1, L2-2, L2-5 and unspecific L2 sublineages (L2-like). **e**, Mosaic composition of hexaploid wheat D genome template tracing back to *Ae. tauschii* sublineages, with L2-1, L2-2 and L2-5 as the top-three donor sublineages. AHGs not able to be assigned with specific sublineages were classified as L1-like, L2-like and L3-like based on the genetic similarities, respectively. **f**, Geographical distribution of *Ae. tauschii* sublineages. The accessions of the top-three donor sublineages, L2-1, L2-2 and L2-5, were colored.

### Evolution trajectories of three *Ae. tauschii* lineages and inter-lineage gene flow

To clarify the lineages delimitation of *Ae. tauschii*, we constructed the phylogenetic tree of *Ae. tauschii* based on the 837 high-confidence mutations in the chloroplast genome, with two *Ae. longissima* accessions as the outgroup (**Supplementary Table 2**). Consistent with previous reports, lineage 1 (L1, *Ae. tauschii* ssp. *tauschii*) and lineage 2 (L2, *Ae. tauschii* ssp. *strangulata*) accessions formed two distinct clades, respectively (**Fig. 1a, Supplementary Fig. 3)**. Six out of nine putative lineage 3 (L3) accessions formed a separate clade from the L1 and L2 clades, indicating the presence of L3, a genetically distinct lineage of *Ae. tauschii*. Consistent with the cytoplasmic phylogeny, there are three distinct clades corresponding to three *Ae. tauschii* lineages in both AHG-based and SNP-based nuclear phylogenies (**Fig. 1a, Supplementary Fig. 4**). Notably, L3 seems to be the basal lineage among the three lineages in both cytoplasmic and nuclear phylogeny, and L2 is the recent lineage. In addition, these six L3 accessions share few ancestries (3.0% on average) with L1 and L2 lineages in the pan-ancestry haploblock map (**Supplementary Fig. 5-6**). The divergence of three *Ae. tauschii* lineages was estimated to have happened about ∼0.2 million years ago (MYA) (**Supplementary Fig. 7**). Based on ABBA-BABA statistics^36^, we detected gene flow from *Ae. longissima* and other Sitopsis D-lineage species to *Ae. tauschii* L1 and L3, indicating that inter-species introgression from D-lineage species might contribute to differentiations of *Ae. tauschii* (**Supplementary Fig. 8**). These results indicate that a total of three lineages of wild goatgrass existed, and L3 is the most ancient gene pool.

**Fig. 3.**
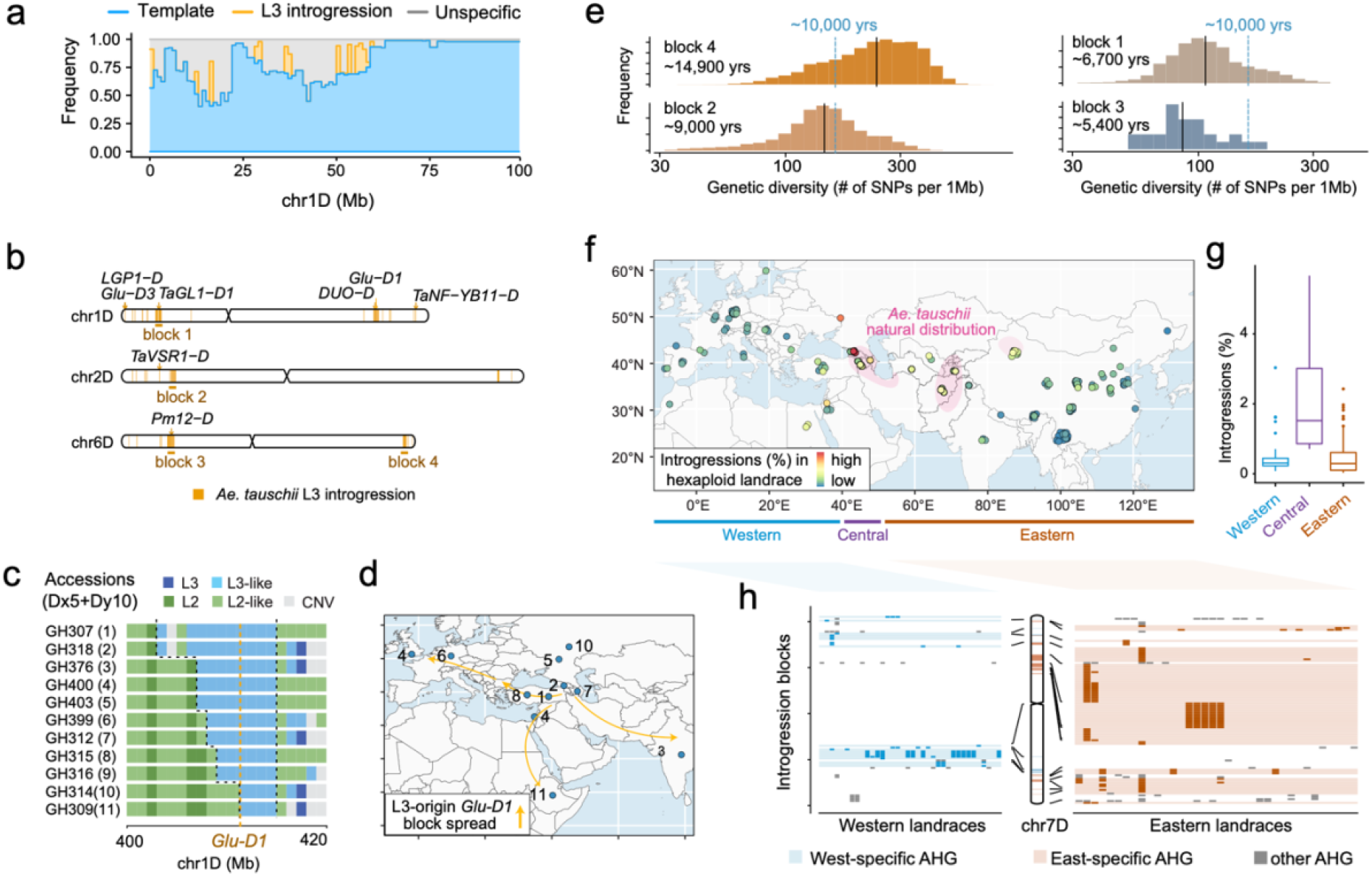
Introgressions from *Ae. tauschii* to bread wheat D genome. **a**, The frequency landscape of L2-origin template AHGs (blue) and L3-origin introgression AHGs (yellow) among hexaploid wheat accessions in the first 100 Mb of chromosome 1D. **b**, The detected *Ae. tauschii* L3-origin introgression regions with low-frequency among hexaploid wheat accessions. Four large introgression blocks (>10Mb) are labeled as block1∼block4. Charaterized adaptive genes are labeled besides the introgression blocks. **c**, Putative origin of *Glu-D1* loci by tracing the ancestral haplotypes across bread wheat accessions with “Dx5+Dy10”. The CDC Stanley assembly was used as the reference genome (x-axis). The boundaries of L3-introgression blocks around *Glu-D1* loci are highlighted with dashed lines. **d**, The geographical distribution of accessions with allele coding “Dx5+Dy10” and the inferred spreading routes of L3-origin *Glu-D1* blocks based on gradual recombination. **e**, Distribution of pairwise genetic distances among hexaploid wheat for four major L3 introgression blocks, which are defined in (**b**). Correspondingly divergence times are estimated by the medians (black line). **f**, Geographical landscape of *Ae. tauschii* introgressions among wheat landraces. Pink circles, the natural distribution area of wild *Ae. tauschii*. **g**, Comparison of the ratios of introgression among hexaploid wheat landraces distributed in western, central and eastern Eurasia, respectively. The geographical ranges of regions are consistent with (**f**). **h**, Presence of introgression AHGs specific to western landraces (blue) and eastern landraces (brown) across chromosome 7D. Accessions are ordered by longitudes. The geographical ranges of regions are consistent with (**f**).

**Fig. 4.**
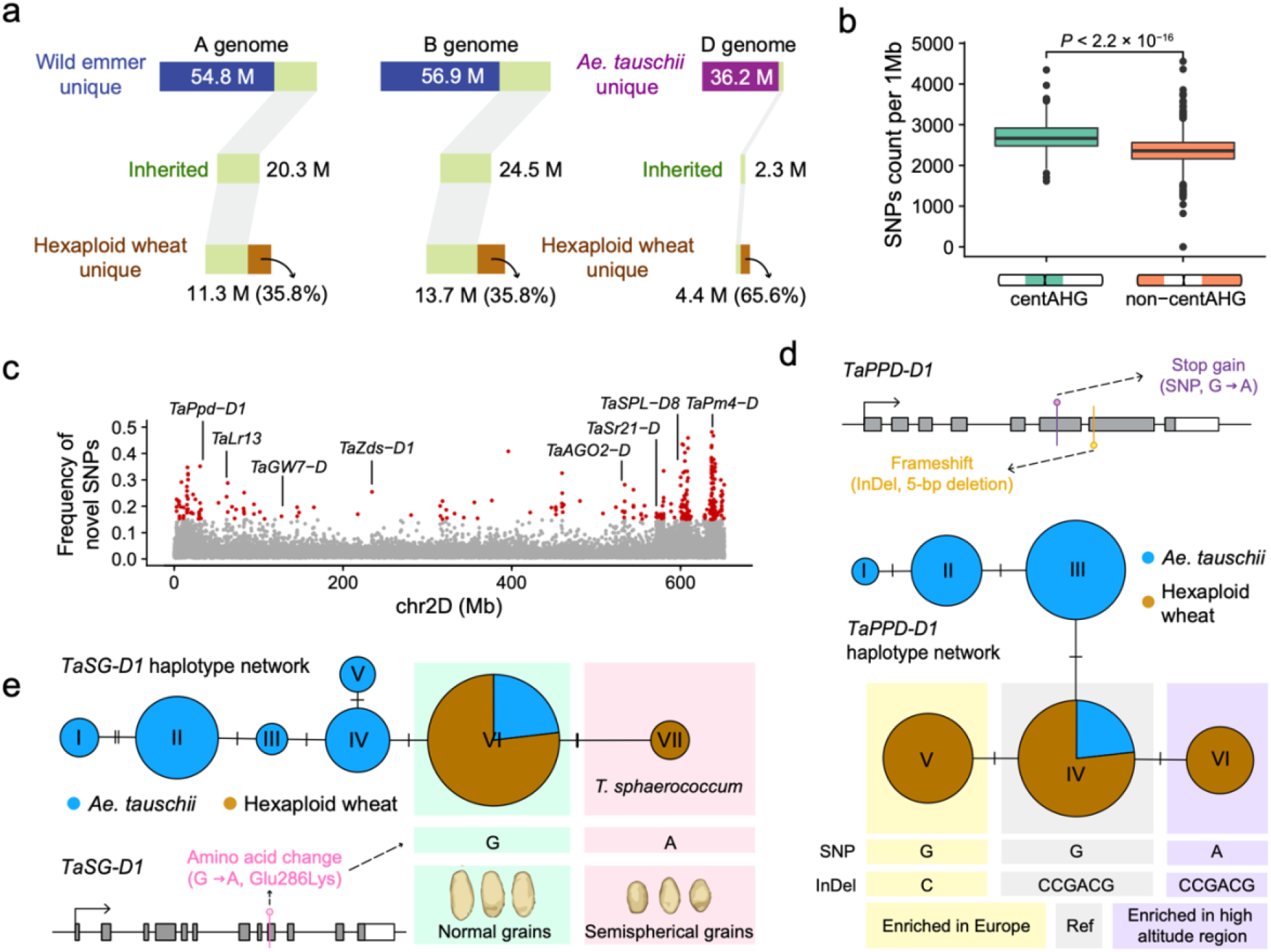
Accumulation of novel mutations in hexaploid wheat D genome. **a**, Schematic diagram shows counts of SNPs unique to the progenitors of hexaploid wheat landrace (A&B genome: wild emmer; D genome: *Ae. tauschii*), counts of SNP inherited by landraces from progenitors (middle) and counts of SNPs unique to hexaploid wheat landrace (bottom). **b**, Comparison of counts of novel SNPs in the centAHG and non-centAHG zones among the D chromosomes (two-tailed *t*-test). **c**, Frequency distribution of novel mutations across chromosome 2D. Hotspots (top 5%) and its flanking regions are highlighted in red. Characterized genes within hotspot regions residing non-synonymous novel mutations are labeled. Bin size, 10-kb. **d**, Haplotype network and gene structure for *TaPPD-D1*, highlighting a novel SNP and a 5-bp deletion found only in hexaploid wheat, leading to stop gain and frameshift, respectively. The A allele of the SNP is enriched in high-altitude accessions^34^, and the 5-bp deletion was enriched in European accessions^47^. Haplotypes were consecutively labeled using Latin numbers according to the evolutionary relationship. **e**, Haplotype network and gene structure for *TaSG-D1*, highlighting a novel SNP leading to an amino acid change, which was only present in hexaploid wheat *T. sphaerococcum*. This SNP was show to be causal for the trait of semispherical grains and improved heat tolerence^48,49^. Haplotypes were consecutively labeled using Latin numbers according to the evolutionary relationship.

**Fig. 5.**
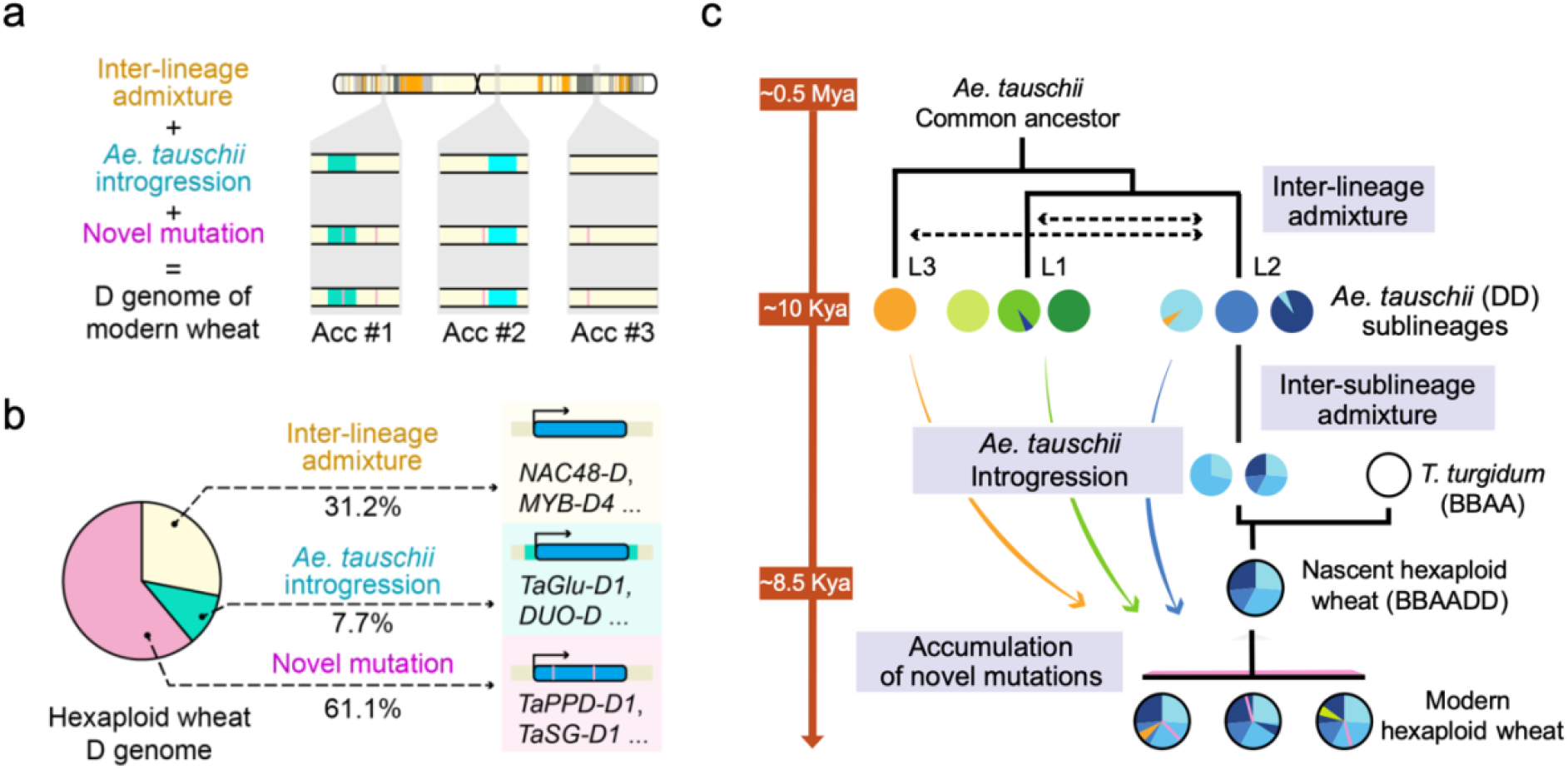
Schematic illustration of the evolution of hexaploid wheat D genome genetic diversity structure. **a**, Schematic diagram shows the three major evolutionary events in the formation of genetic diversity of hexaploid wheat D genome. Hexaploid wheat shared the same set of ancestral mosaic as the genomic template resulted from the admixture of *Ae. tauschii* sublineages (inter-lineage admixture). After hexaploidization, large genomic blocks introgressed to hexaploid wheat from *Ae. tauschii* lineages. Novel mutations accumulated and further shaped the genetic diversity of D genome (novel mutation). **b**, Proportion of hexaploid wheat D genome segments dominantly shaped by three classes of evolutionary approaches, with corresponding genes of each class labeled. *Ae. tauschii* introgression, identified *Ae. tauschii* introgressions. Novel mutation, genomic regions with novel mutation rate > 1/1000. Inter-lineage admixture, the remaining regions. **c**, Genomic evidence informed evolutionary model of hexaploid wheat D subgenome (right), and the corresponding time scale (left). Green, blue and orange sectors denoting the *Ae. tauschii* L1, L2 and L3 origin genetic diversity, respectively. The types of AHGs were discriminated by their brightness. Succinct descriptions of four major event involving in the evolution of hexaploid wheat D subgenome are denoted with blue shading. Arrow colors indicate the phylogenetic relatedness between taxa. Black, direct pedigree. Colored, genetic introgression. This schema is based on the results from this study and prior assumptions from the literature.

As indicated by both cytoplasmic and nuclear phylogenies, three putative L3 accessions (AE041, AE063 and AE117) are clustered within the L1 clade, which contrasts their reported subspecies delimitation (**Fig. 1a**). We found in the three accessions, up to 98.3% genomic segments can be traced to the ancestral haplotypes shared with *Ae. tauschii* L1 or L2 (**Fig. 1b**), and most segments traced to L1 or L2 were clearly demarcated without overlaps, indicating their cross-lineage hybridization origin (**Fig. 1c, Supplementary Fig. 9**). Further clustering analysis confirmed that the three putative L3 accessions have hybrid ancestry from L1 and L2, while the ancestries of other six accessions were specific to L3 (**Supplementary Fig. 10**). These three accessions were labeled as hybrids and excluded from following analyzes (**Supplementary Table 1**). These findings confirmed the existence of genetically distinct L3 and suggested that some putative L3 accessions are sublineage intermediates that may originate from hybridization events.

To provide an overview of gene flow among *Ae. tauschii* lineages and further clarify population structure, we performed clustering analysis and constructed the phylogenetic network to evaluate levels of admixture among lineages. By estimating the ancestry coefficients of each accession, we identified 148 intermediate accessions among sublineages, which counts nearly one-third of the total *Ae. tauschii* accessions (**Fig. 1d**). Consistent with the phylogenetic tree, accessions from *Ae. tauschii* L1 and L2 formed two major branches in phylogenetic networks, and many splits existed among sublineages, indicating the existence of reticulate events among sublineages (**Supplementary Fig. 11**). These results suggested that gene flow occurred among *Ae. tauschii* sublineages. By selecting accessions with major ancestral coefficients > 99%, 225 non-admixture *Ae. tauschii* accessions are further clustered into 13 sublineages based on genomic ancestries and geographical distribution (**Fig. 1e, Supplementary Table 3**). Taken together, our results suggest that, *Ae. tauschii* were composed of genetically distinct sublineages and the inter-sublineage geneflow played a key role in shaping the diversity of *Ae. tauschii* (**Fig. 1f**).

### Unified ancestral template of hexaploid wheat D genome with mosaic origins

Based on the refined population structure of *Ae. tauschii*, we further investigated the evolutionary process leading to the origin of the hexaploid wheat D genome. Principle coordinate analysis (PCoA) showed that hexaploid wheat accessions were grouped in a narrow clade, consistent with the limited genetic diversity compared with *Ae. tauschii* (**Fig. 2a**). In 59.6% of D genomic regions, there was only one AHG type across all hexaploid wheat accessions, in contrast to a median of 22 AHG types in *Ae. tauschii* (**Fig. 2b**). These results indicated a conserved AHG composition of hexaploid wheat D subgenome presenting as a genetic diversity “template”. All the hexaploid wheat accessions shared this genomic template with few AHGs specific to each (sub-)species (**Supplementary Fig. 12, Supplementary Table 4**). We further identified D genomic regions covered by the conserved template where the frequency of dominant AHG exceeded 90% in hexaploid wheat, and the dominant AHG of each genomic window constituted the template (**Supplementary Fig. 13, Supplementary Table 5**). This genomic template covers 88.5% of D genomes, especially centromeric regions of each chromosome. 4D was the chromosome with the highest template coverage (98.0%) among the seven D chromosomes, while large genomic blocks outside the template existed on the distal region of chromosomes 1D and 2D. We show that the genetic diversity of the hexaploid wheat D genome had been fixed at the dawn of the hexaploid wheat evolution, as divergence levels are similar across the backbone region (**Supplementary Fig. 14**).

To investigate the evolutionary process leading to the genomic template of the hexaploid wheat D genome, we traced AHGs from the hexaploid wheat D genome back to *Ae. tauschii*, and 84.9% of AHGs could be traced back to collected *Ae. tauschii* accessions (**Fig. 2a, Supplementary Table 4**). Though dominant AHGs in other genomic regions, including the central region of 3D (181 - 338 Mb) and 7D (264 - 387 Mb) chromosomes, could not traced to specific *Ae. tauschii* sublineages, we confirmed their *Ae. tauschii* L2 origin (**Supplementary Fig. 15 - 16, Supplementary Table 6**). At the genome-wide scale, the genomic template showed a mosaic pattern of ancestries mainly derived from L2 sublineages (**Fig. 2d, 2e**). Genome-wide AHG-based phylogeny confirmed *Ae. tauschii* L2 as the direct donor of hexaploid wheat D subgenome, as in previous reports^21^ (**Supplementary Fig. 17**). The admixture level of the genomic template is much higher than that observed among *Ae. tauschii* accessions **(Supplementary Fig. 18)**.

Among the *Ae. tauschii* sublineages, the L2-2 sublineage contributed most AHGs (49.2%) to the hexaploid wheat D subgenome template, suggesting their role as major donors in the early formation of the hexaploid wheat D genome. Accessions from this sublineage were mainly distributed southwest of the Caspian Sea (Fig. 2f), where the hexaploidization event was proposed to happen^21^. Sublineage L2-5, mainly distributed east of the Caspian Sea, contributed 20.3% of the template. Up to 7.0% of the wheat D genome template could be traced back to *Ae. tauschii* L2-1 and accessions from this sublineage were mainly distributed around the southeast of the Caspian Sea. The Southern Caspian coastal region has been reported as the genetic diversity center of *Ae. tauschii*^12^. These three sublineages contributing most to the template were estimated to differentiate approximately 70,000 ∼ 90,000 years ago, earlier than hexaploidization was estimated to happen **(Supplementary Fig. 7)**. Notably, an L3-origin AHG was also fixed as a part of the template (7D:81-82 Mb), which might result from the admixture of *Ae. tauschii* sublineages before the hexaploidization.

By quantifying genetic diversity depletions of hexaploid wheat compared with *Ae. tauschii* across the D genome, we found a ∼10Mb segment on the short arm of chromosome 2D among regions with the highest selection intensities **(Supplementary Fig. 19)**. This region overlapped with the fine mapping interval of *TaTg-2D*, vital in regulating the hullness of the spike, one of wheat domestication syndrome^37^. A subset of this region (23-24Mb) was almost fixed in hexaploid wheat, and the dominant AHG of this region could be traced back to *Ae. tauschii* L2-2 **(Supplementary Fig. 19)**. Another key wheat domestication gene *BTR-D* was assumed to be involved in the change of rachis brittleness after the formation of hexaploid wheat^32,38^. The genomic block residing homologs of *BTR-D* also showed a severe reduction of diversity, within which we identified mutations that were diversified between wild goatgrass and hexaploid wheat **(Supplementary Fig. 20)**. These results show that the hexaploid wheat shared the same set of ancestral mosaic genomic template across the D genome, which originated mainly from an admixture of three of *Ae. tauschii* sublineages.

### *Ae. tauschii* introgression events reshaped the bread wheat D genome

To investigate the genetic influence of introgression from *Ae. tauschii* to bread wheat D subgenome, we systematically identified alternative AHGs besides the dominant AHGs composing the template, especially those that could be traced back to *Ae. tauschii* (**Fig 3a, Supplementary Table 7)**. These alternative AHGs existed at relatively low frequencies in the hexaploid wheat population (1.9% on average), in contrast to the template AHG (98.3% on average) **(Supplementary Fig. 21)**. ABBA-BABA statistics^36^ and topology weighting approach^39^ further confirmed their putative introgression origin (**Supplementary Fig. 22 - 23**). We profiled the total length of all alternative AHGs in each hexaploid wheat accession **(Supplementary Fig. 24)**. Chinese cultivar ChuanMai42 preserved the highest ratio of alternative AHGs, its 625 Mb continuous alien introgressions were mainly distributed on chromosome 3D and 4D (**Supplementary Fig. 25**). Considering ChuanMai42 was derived from recent introgression breeding^40,41^, it was excluded from further analysis of introgressions. Few introgressions were found in hexaploid wheat landraces from Yunnan province of China, probably due to their isolated geographical distribution.

Among the three lineages, the L3 lineage contributed the most introgressions to the hexaploid wheat D genome, for 7.4 Mb on average in each accession. L2 lineage contributed 4.3 Mb on average in each accession (**Supplementary Fig. 26**), and L1 origin introgression contributed 1.7 Mb on average per accession, which is the least among the three lineages of *Ae. tauschii* (**Supplementary Fig. 27)**. Notably, 13 Mb of L3-origin AHG introgression blocks in hexaploid wheat also presented in L2 accessions, indicating that some of the L3 introgression was first introgressed into L2 through lineage admixture and further combined into the hexaploid gene pool (**Supplementary Fig. 28**). Four major L3 introgression blocks were found on chromosomes 1D, 2D, and 6D, respectively **(Fig. 3b)**. The length of these four blocks ranges from 7 Mb (chr6D: 457 - 464 Mb) to 10 Mb (chr6D: 71 - 81 Mb).

Genes underlying key agronomic traits like *Glu-D3*^24^ and *Pm21*^42^ resided in the L3 introgression region (**Supplementary Fig. 29 - 30**). Compared with tetraploid wheat, hexaploid wheat outperforms for the superior baking quality, in which trait HMW glutenin in the D subgenome played an important role, and the HMW-GS “Dx5+Dy10” has shown positive effects on dough properties^43,44^. We found that all 14 collected “Dx5+Dy10” accessions show a distinct pattern of normalized read depth for the 1 Mb genomic window residing *Glu-D1* loci, compared with “Dx2+Dy12” accessions **(Supplementary Fig. 31)**. To trace the origin of allele coding “Dx5+Dy10”, we changed the reference genome from Chinese Spring (“Dx2+Dy12”) to CDC Stanley (“Dx5+Dy10”). We found the genomic window residing *Glu-D1* was of L3 origin **(Fig. 3c)**, indicating the superior allele coding “Dx5+Dy10” was introgressed from the *Ae. tauschii* L3 to the hexaploid wheat^22^. The introgression block residing *Glu-D1* had been gradually broken by recombination along with the spreading of bread wheat from the Middle East **(Fig. 3d)**. These analyses demonstrated the important role of introgression from lineages of *Ae. tauschii* in the genetic diversity of bread wheat D genome.

To provide a spatial-temporal overview of *Ae. tauschii* introgression, we further quantified the diversification of four major L3 introgression blocks among hexaploid wheat to date their occurrences. The result showed that the most ancient introgression event (block 4) occurred before hexaploidization at ∼14,900 years before present (YBP) **(Fig. 3e)**. Introgression into hexaploid wheat gene pool continuously occurred after hexaploidization from ∼9,000 YBP (block 3) to ∼5,400 YBP (block 2). Ratios of introgression blocks in the genome vary among hexaploid wheat accessions. Landraces with a relatively high proportion of introgressions were located around the Transcaucasia region, where *Ae. tauschii* L2 was naturally distributed (**Fig. 3f**). The introgression ratios declined westwards and eastwards from the Transcaucasia region, indicating the gradual loss of introgression during spreading (**Fig. 3g**). 34.3% of total introgression AHGs are specific to western or eastern landraces, and most introgression AHGs were first shown in the near west and east accessions along spreading **(Fig. 3h)**. This pattern was not found in the hexaploid wheat cultivar (**Supplementary Fig. 32**). Together, our results suggest that, large genomic blocks from *Ae. tauschii* had been introduced into the hexaploid wheat D genome, frequencies of which declined along with independent western and eastern spreading from the origin of hexaploid wheat in the Middle East.

### Novel mutations accumulated in the hexaploid wheat D genome during the spreading

Wheat compensates for genetic bottlenecks and adaptation to various environments partially by generating new diversity at a relatively fast pace^4,45^, while the genetic contribution of novel mutations on the hexaploid wheat D genome remains unclear. By excluding SNPs identified in both hexaploid wheat landrace D genome and *Ae. tauschii*, we found that nearly two-thirds (65.6%) of the SNPs in the hexaploid wheat D genome are novel mutations **(Fig. 4a)**. In contrast, only about one-third (35.8%) SNPs in hexaploid wheat A&B subgenome could be found in wild emmer wheat, the tetraploid progenitor. Only a minority of novel mutations were preserved in individual varieties, and population-wide frequencies of novel mutations were relatively low compared with inherited mutations **(Supplementary Fig. 33)**. The contrast in the ratio between D and A&B genomes is partially due to the pervasive interploidy introgression from tetraploid wheat to hexaploid wheat, while the introgression from *Ae. tauschii* to hexaploid hardly happened. This contrast highlights the important role of novel mutations in the formation of genetic diversity in the hexaploid wheat D genome.

The density of novel mutations displayed clear patterns at both chromosome-wide and local scales **(Supplementary Fig. 34)**. Counts of novel mutations vary significantly among local regions, and a total of 729 novel mutations hotspot (top 5%) genomic regions were identified, extending 174.2 Mb in length (**Supplementary Table 8**). Chromosome 4D harbors most hotspot regions (44.8 Mb), while chromosome 1D harbors the least (13 Mb). There are more novel mutations in the centric regions than in the distal regions (one-tailed *t*-test, *P* < 2.2 × 10^−22^) **(Fig. 4b)**.

Some novel mutations with relatively high frequencies were annotated as non-synonymous mutations **(Fig. 4c)**. We identified a novel SNP and a novel InDel leading to the stop gain and frameshift of *TaPPD-D1* **(Fig. 4d)**, which gene is the major determining factor affecting the photoperiod response and vital in the broad adaptation of hexaploid wheat^46^. The A allele of the SNP was enriched in high-altitude accessions^34^, while the 5-bp deletion was specifically utilized in European wheat germplasms^47^. An SNP in *TaSG-D1* was confirmed to be causal for the taxonomically recognized trait, semispherical grains, and improved heat tolerance for hexaploid wheat subspecies *T. sphaerococcum*^48,49^, and we confirmed that this SNP was also a novel SNP absent in *Ae. tauschii* **(Fig. 4e)**. Novel mutations were also identified in key genes regulating key agronomic traits, such as in *TaGS5-D1* regulating grain size and in *TaGW7-D* regulating grain width **(Supplementary Fig. 35 - 36)**. These results demonstrated novel mutations impacted the formation of genetic diversity of the hexaploid wheat D genome, and also the diversification and adaptation of hexaploid wheat.

### Reconcile the genetic diversity of hexaploid wheat D genome in evolutionary view

Our results revealed a hierarchical architecture of genetic diversity in the bread wheat D genome **(Fig. 5a, Supplementary Fig. 37)**. The ancestral mosaic genomic template originated mainly from the inter-lineage admixture among *Ae. tauschii* sublineages, composing the first layer of bread wheat D genome genetic diversity. Introgressions from *Ae. tauschii* into the hexaploidization gene pool partially replaced the genomic template. Along the evolution, novel mutations continuously accumulated in the hexaploid wheat D genome **(Fig. 5a)**. These three evolutionary forces played important roles in the formation of the extant genetic diversity, as well as the genes in hexaploid wheat D genome **(Fig. 5b)**. Collectively, 7.7% of hexaploid wheat D genomic regions were among *Ae. tauschii* introgressions and 61.1% of genomic regions resided in novel mutations. The other 31.2% of genomic regions remained generally unchanged as the genomic template.

According to the hierarchical structure of genetic diversity in the hexaploid wheat D genome, we proposed a refined evolutionary model of the hexaploid wheat D genome, involving recurring episodes of hybridization and gene flow **(Fig. 5c)**. Over a long time since its origin, *Ae. tauschii* (DD) differentiated into lineages, among which high genetic diversity accumulated. Later, genetic admixture happened among lineages *Ae. tauschii*. During the Neolithic period (∼10,000 years ago), hexaploid wheat emerged between domesticated tetraploid wheat and the admixture of *Ae. tauschii* mainly from three sublineages distributed among the southwestern Caspian Sea region, with two possible evolutionary scenarios **(Supplementary Fig. 38)**. The additional D genome in hexaploid wheat introduced severe genomic diversity reduction and fixed the genomic template in hexaploid wheat, supporting its further spread than tetraploid wheat. After hexaploidization, large introgression genomic blocks from *Ae. tauschii* were introduced and gradually lost along with the independent western and eastern spreading of hexaploid wheat from the Middle East. Along the evolution of nascent hexaploid wheat, novel mutations accumulated and further shaped the modern hexaploid wheat D genome.

## Discussion

There has been a seeming incongruity in the genetic diversity of the hexaploid wheat D genome for a long time, as it has been proposed that a few *Ae. tauschii* participated in the formation of bread wheat, while evidence supporting the multiple hexaploidization accumulated^13,14,21^. By leveraging the pan-ancestry *Triticum-Aegilops* D genome haploblock map constructed based on 762 genomes of *Ae. tauschii* and hexaploid wheat, we revisited the evidence for the origin of the hexaploid wheat D genome and its evolutionary relationship with *Ae. tauschii*.

### Inter- and intra-specific introgression contributed to the evolution of *Ae. tauschii*

Our results suggest that gene flows occurred among three genetically distinct sublineages of *Ae. tauschii*, and independent intermediate accessions among sublineages were identified, implying that despite being a self-pollinating species, intra-specific introgression contributed to the evolution of *Ae. tauschii*^50,51^. Consistent, remnants of wild goatgrass have been identified at several pre-agricultural settlements, and co-cultivation as a weed over an extended period might result in the mixing of *Ae. tauschii* from different geographical origins^52^. We further detected introgression from Sitopsis species of D-lineage *Aegilops* to *Ae. tauschii*, indicating that inter-species introgression contributed to its differentiations, though the possibility of incomplete lineage sorting could not be fully ruled out.

### The formation of ancestral mosaics in the bread wheat genome

It has been proposed that the limited genetic diversity of the hexaploid wheat D genome resulted from the hexaploidy genetic bottleneck, and only a small population of *A. tauschii* contributed to the formation of bread wheat^20,35^. However, highly diversified genomic regions existed between the hexaploid wheat D genome and each of the assembled *Ae. tauschii* genomes^13,21^. This incongruity further raised the question of how nascent hexaploids with limited genetic diversity adapted to diverse environments. We showed that all hexaploid wheat, including all hexaploid wheat (sub-)species^53^, shared a conserved genomic template across the D genome, the mosaic composition of which was mainly an admixture of three *Ae. tauschii* L2 sublineages, which could reconcile the seeming incongruity. We proposed two potential models for the forming of such a mosaic template. One is that the hexaploidization happened between tetraploid wheat and a high admixture *Ae. tauschii* accession, which is less likely considering the level of admixture among *Ae. tauschii* is much lower than that observed in the genomic template. The other is that hexaploidization happened multiple times between tetraploid wheat and various *Ae. tauschii* accessions and only one combination template thrived and replaced others.

Together with our previous work^33^, we demonstrated that all three subgenomes of hexaploid wheat showed mosaic ancestral patterns. This mosaic genome combined genetic diversity from various sources and enabled the pyramiding of the superior alleles^4^, leading to improved adaptability and the evolutionary success of bread wheat. Nevertheless, the mosaic patterns of the D genome are quite different from the A&B genomes among individual varieties, as the D genome of bread wheat keeps a unified mosaic pattern, in contrast to the diverse ancestry composition observed in A&B genomes. This contrast was partially due to the following two reasons. One is that the beginning of tetraploid wheat domestication was ∼3,000 years earlier than the origin of hexaploid wheat^32^, during which multiple templates of diverse ancestry composition appeared in the tetraploid wheat through pervasive hybridizations under early agricultural activity, which were further inherited by hexaploid wheat. Two is the probability difference of natural gene flow from *T. turgidum* and *Ae. tauschii* to hexaploid wheat. Differences in ploidy presented only a weak barrier to gene flow from tetraploid wheat to hexaploid wheat, leading to the introgression of diverse ancestries into A&B genomes, while hybridization between *Ae. tauschii* and hexaploid wheat rarely occurred in nature^54^.

### Multi-layered variations accumulated in the D genome

The inclusion of the D genome is important to the board adaption and superior end-use quality for hexaploid wheat^4,55^. Due to the unified genomic template, hexaploid wheat only inherited limited genetic diversity from *Ae. tauschii* in the D genome, lower than that of inherited from tetraploid wheat in A&B genomes. Therefore, the relative importance of introgression and accumulation of novel mutation is much higher for the genetic diversity of the D genome. Considering the rare natural hybridization between *Ae. tauschii* and hexaploid wheat, tetraploid wheat may have served as a bridge for gene flow from *Ae. tauschii* into hexaploid wheat, and introgression is more likely to arise from the areas where swarms of both tetraploid wheat and hexaploid wheat existed^11,21^. In addition, it needs to be mentioned that the inferred novel mutations could be somehow biased, considering the collected *Ae. tauschii* accessions are still limited and may be different from those at the dawn of hexaploidization. The accumulation of variations caused the multi-layered genetic diversity structure of the D subgenome. Many important genetic loci such as *TaBTR-D1, TaPPD-D1*, and *TaGlu-D1* gained genetic diversity through this featured structure, which ensured adaptation and agronomic utilization of hexaploid wheat.

### Innovation in the D genome architecture brings success to the bread wheat

The mosaic origin has been reported in several plant species, such as maize^56^, Einkorn wheat^57^, oat^58^, barley^59^, etc, that genetic variation was contributed across a geographic mosaic of populations^60^. At the early stage of hexaploidization, there might been multiple ancestral compositions in nascent hexaploid wheat. These nascent hexaploids with various transient ancestral blocks combination hybridized with each other for many rounds and underwent selection as a genetically compound species continuum before a dominant genotype with the genomic template we observed appeared^61^. This situation possibly resulted from early agricultural activity^62^, the possibility of inter-lineage hybridization arose with the company of human behavior, and early farmers selected combinations fitting early agricultural needs. This indicates that the AHG template might be an optimized one, which was selected from a pool of mosaic templates by rare chance under early agricultural activity. This also inspired that for *de novo* domestication, creating and selecting novel mosaic D genome templates, rather than using current *Ae. tauschii* germplasms directly in synthesizing hexaploid wheat may accelerate the process for pyramid beneficial alleles.

The addition of the D genome rendered hexaploid wheat with high grain yield and superior quality, and supports its further spreading exceeding the geographical distribution of tetraploid wheat. Our result indicated the demography has largely shaped the genomic variation of the hexaploid wheat D genome^63^. For future wheat breeding, the identified alternative haplotypes in the hexaploid wheat D genome could be utilized directly, and our results also highlight the importance of crop wild relatives as their potential as a source of adaptive diversity.

