## Supplementary figures for "Tracing the genetic diversity of the bread wheat D genome"

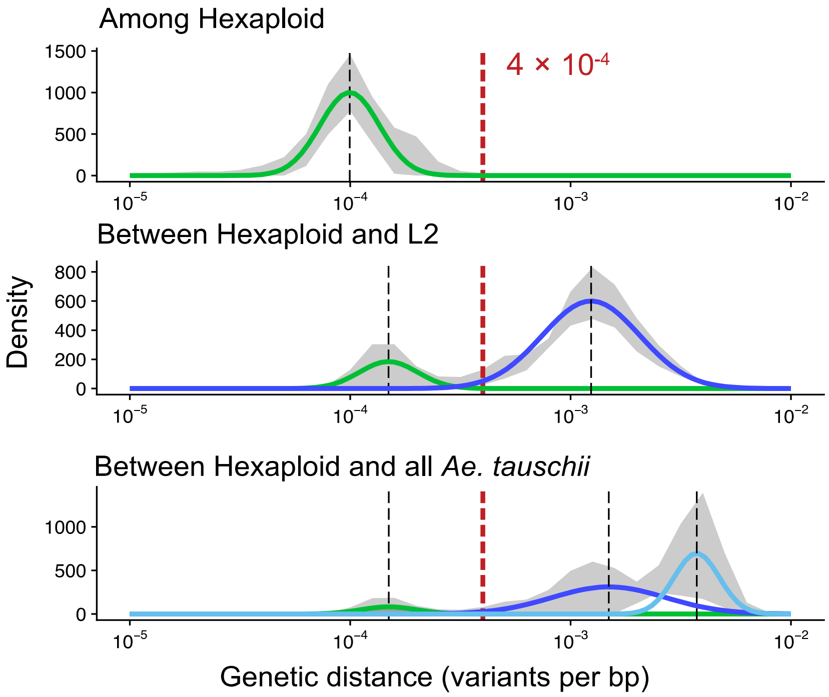


**Supplementary Fig. 1 Distribution of pairwise genetic distances for three sample pair combinations in the D genome.** Upper: among hexaploid wheat. Middle: between hexaploid wheat and *Ae. tauschii* L2. Bottom: between hexaploid wheat and all *Ae. tauschii*. Shadow ribbons, mean ± sd. Solid lines represent the Gaussian distributions fitted by the EM algorithm from 1000 randomly chosen sample pairs in each track. Mean ± sd of the fitted Gaussian distribution are indicated by dashed lines and shadow boxes, with corresponding divergence times labelled. Red line, the threshold at $10^{-3}$ variants per bp.

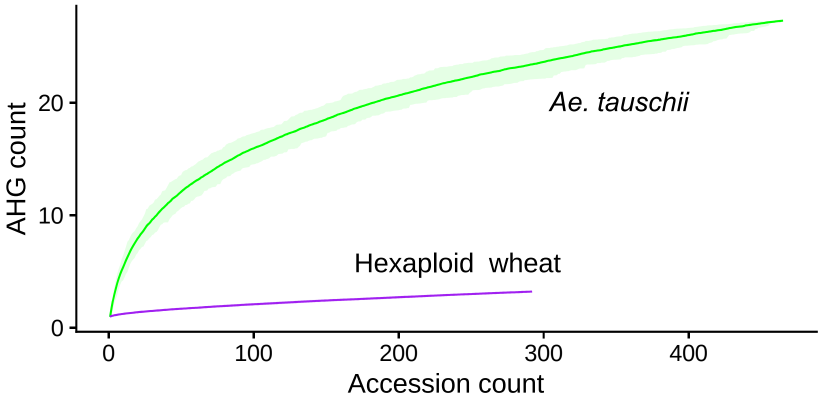


**Supplementary Fig. 2 Saturation curves showing the cumulative number of AHG types versus the number of accessions included.** Solid line and shaded region, average and 90% confidence intervals for 7 D chromosomes.

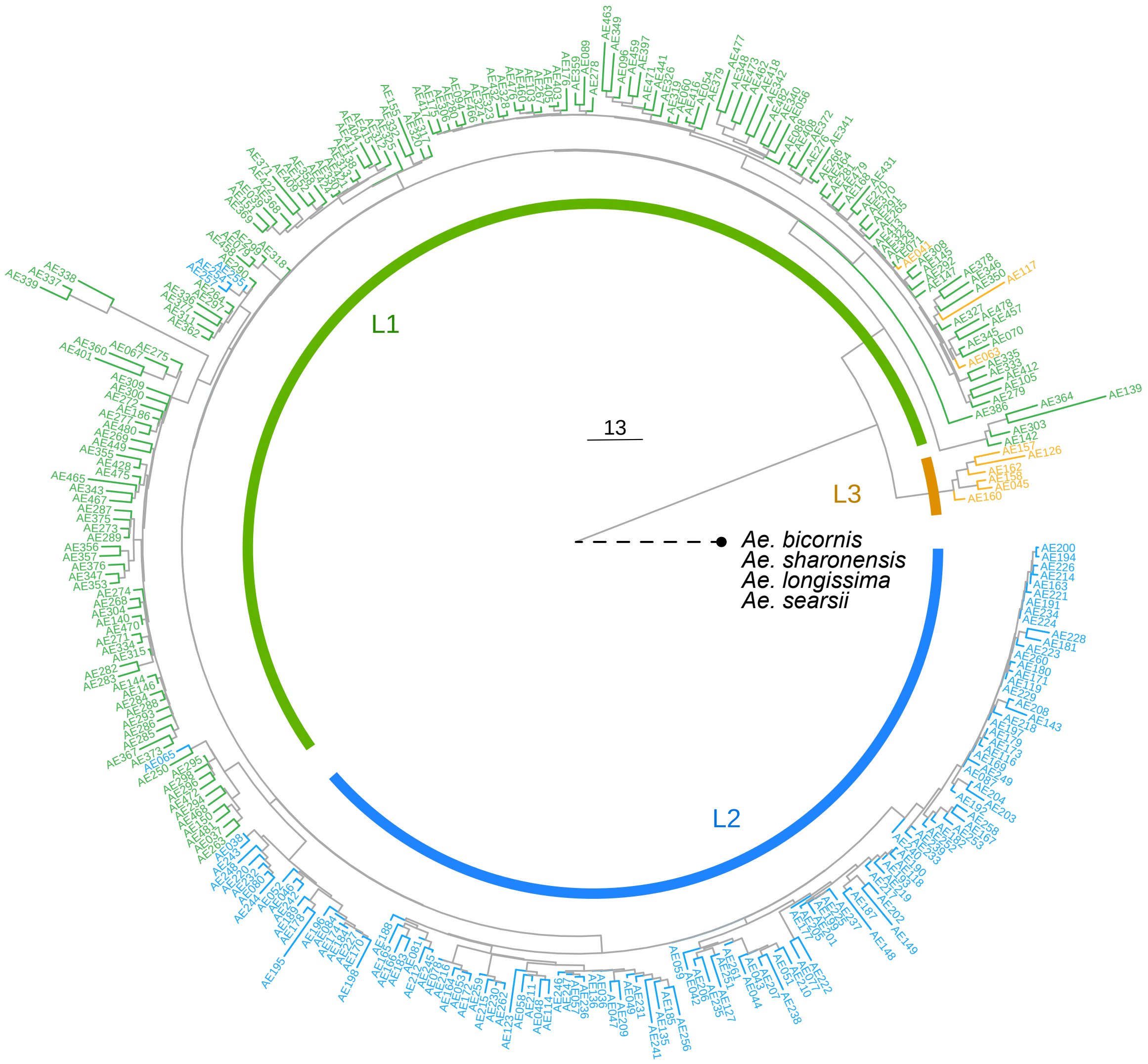


**Supplementary Fig. 3 The chloroplast SNP-based neighbor-joining tree of *Ae. tauschii*.** The chloroplast tree is rooted by assigning *Ae. bicornis*, *Ae. speltoides*, *Ae, longissima* and *Ae. searsii* as the outgroup. Sites with heterozygosity < 0.1 and missing rate < 0.1 were kept for tree construction. The resequencing data of the *Ae. searsii*, *Ae. bicornis*, *Ae. sharonensis* were from (Li et al., 2022)

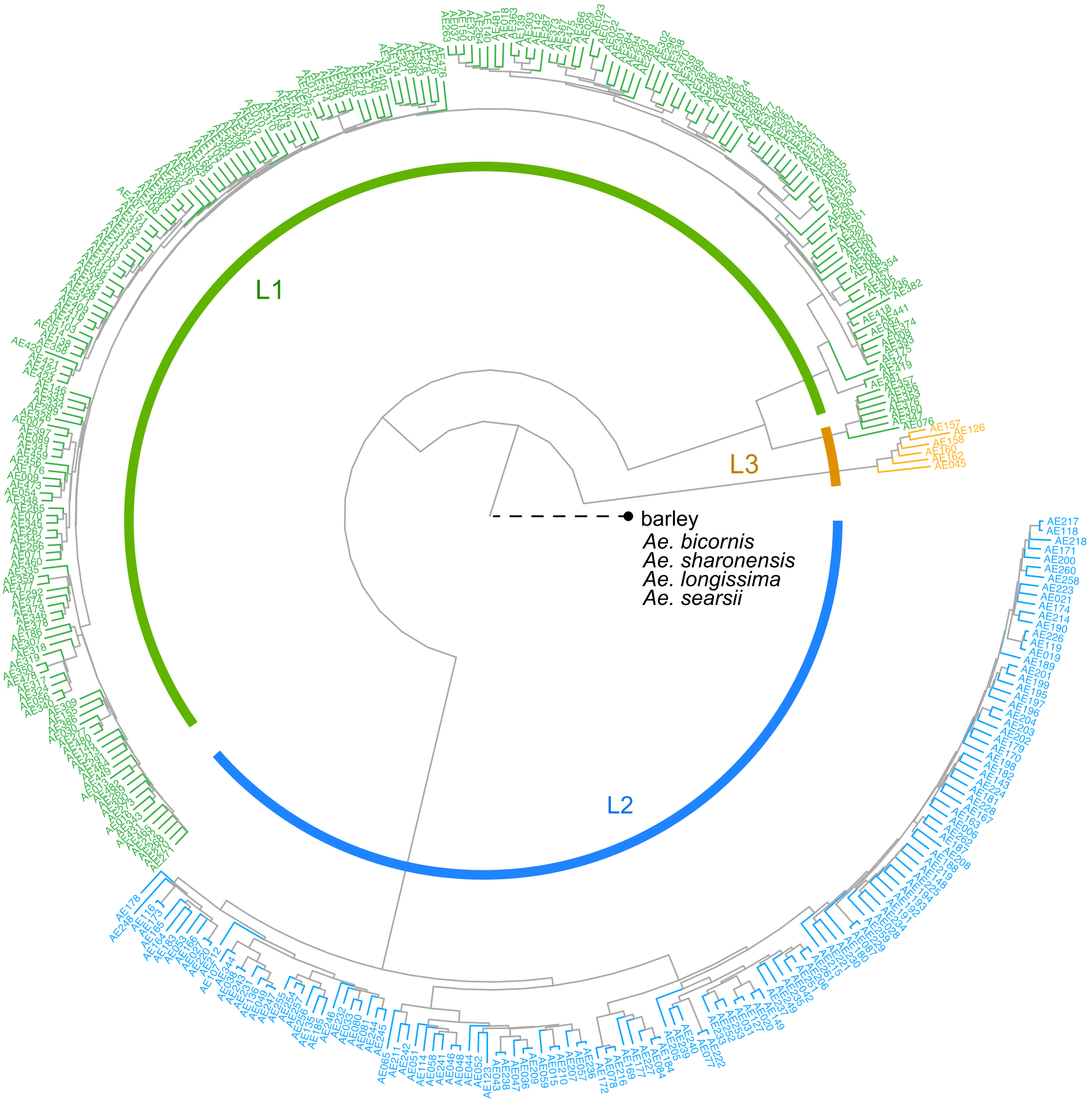


**Supplementary Fig. 4 The D genome SNP-based neighbor-joining tree of *Ae. tauschii***. The chloroplast tree is rooted by assigning barley as the outgroup. Sites with either barley*, Ae. bicornis*, *Ae. speltoides*, *Ae, longissima* or *Ae. searsii* genotyped were kept for tree construction. The resequencing data of the *Ae. searsii*, *Ae. bicornis*, *Ae. sharonensis* were downloaded from (Li et al., 2022).

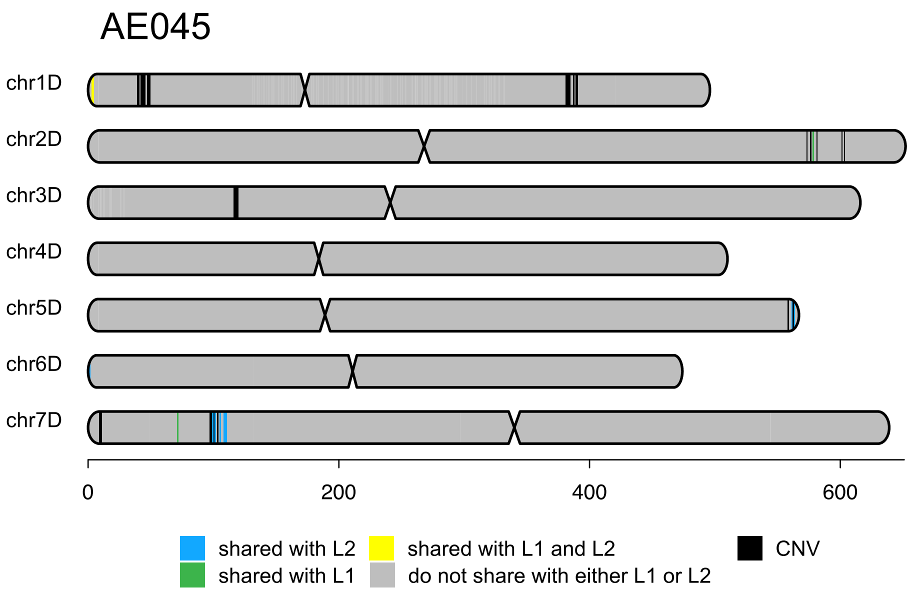


**Supplementary Fig. 5 Chromosomal distribution of ancestry for the putative *Ae. tauschii* L3 accession AE045.**

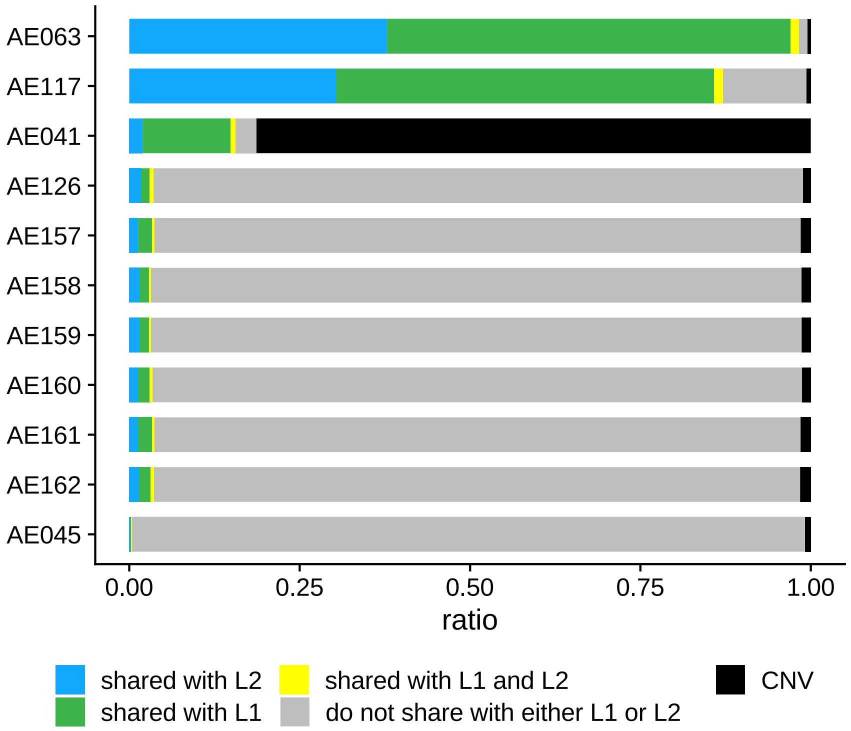


**Supplementary Fig. 6 Ancestry composition of all putative L3 *Ae. tauschii* accessions.** Three intermediate lineage accessions, AE063, AE117 and AE041 show a high ratio of ancestry from *Ae. tauschii* L1 and L2.

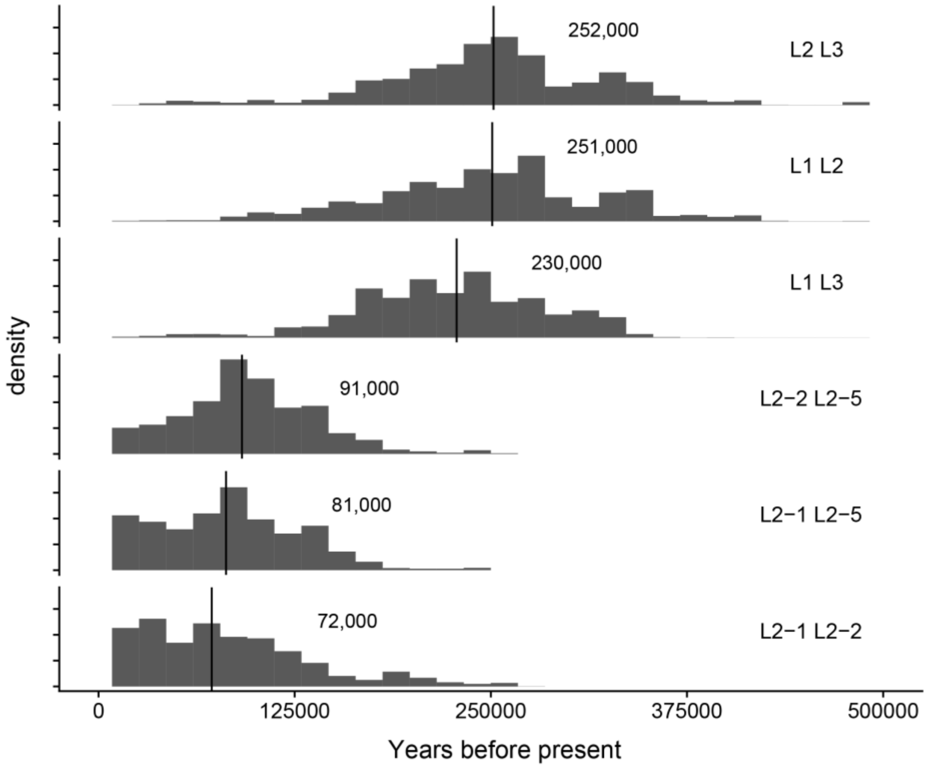


**Supplementary Fig. 7 Distribution of pairwise genetic distances among three *Ae. tauschii* (sub-)lineages across D chromosomes**. L1, L2, L3, three *Ae. tauschii* lineages. L2-1, L2-2, L2-5, three *Ae. tauschii* L2 sublineages. Correspondingly divergence times are estimated by the medians (black line).

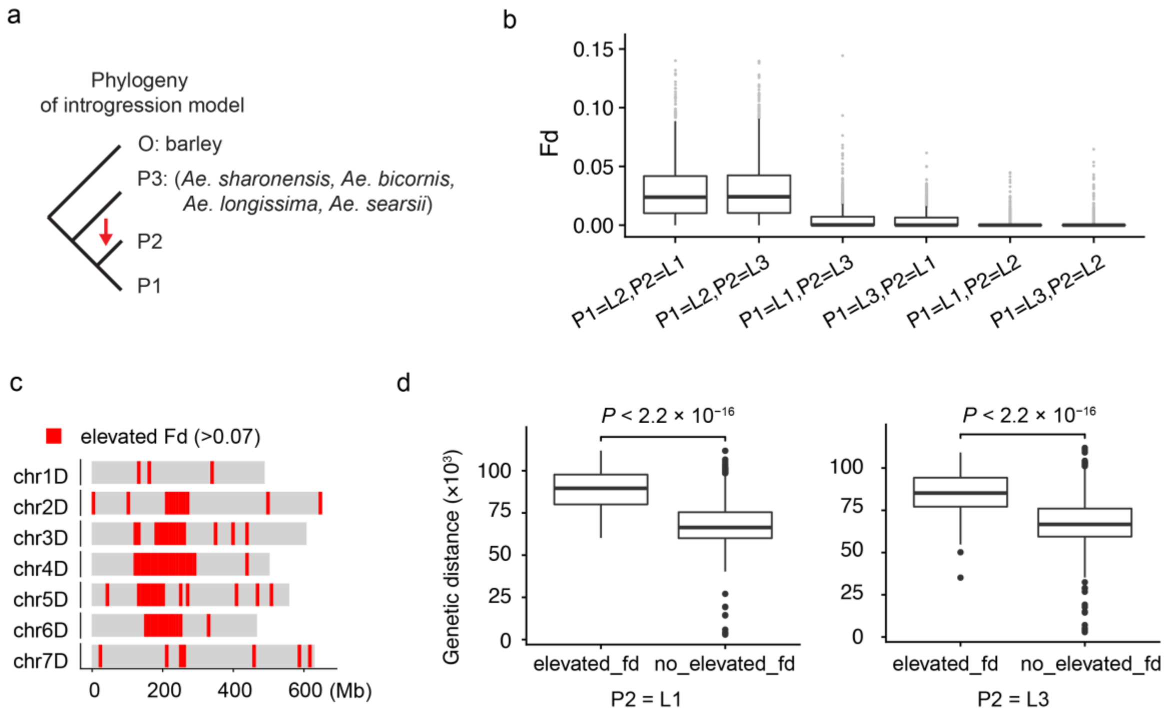


**Supplementary Fig. 8 Detection of introgression from four *Aegilops* section Sitopsis species to three *Ae. tauschii* lineages.** a, The four-taxon topology used for modelling introgression in $f_D$ statistics across D chromosomes. The four terms of each label separated by a comma denotes P1, P2, P3 and O, respectively. L1, *Ae. tauschii* subsp. *tauschii* (L1). L2, *Ae. tauschii* subsp. *strangulata* (L2). L3, *Ae. tauschii* L3. b, Testing gene flow from donor populations to *Aegilops* lineages. gene flow (as indicated by fd) from a specific donor is significantly higher than from wild einkorn. c, The genomic distribution of 10-Mb windows with elevated $f_D$ values ($f_D$ > 0.07) using any of the four Sitopsis species as P3, L1 and L3 as P2. d, Branch length tests showed that the divergence level between *Ae. tauschii* and donor species are significantly higher for genomic regions with elevated $f_D$ values than those with flat $f_D$ values, which indicated introgression rather than incomplete lineage sorting (ILS).

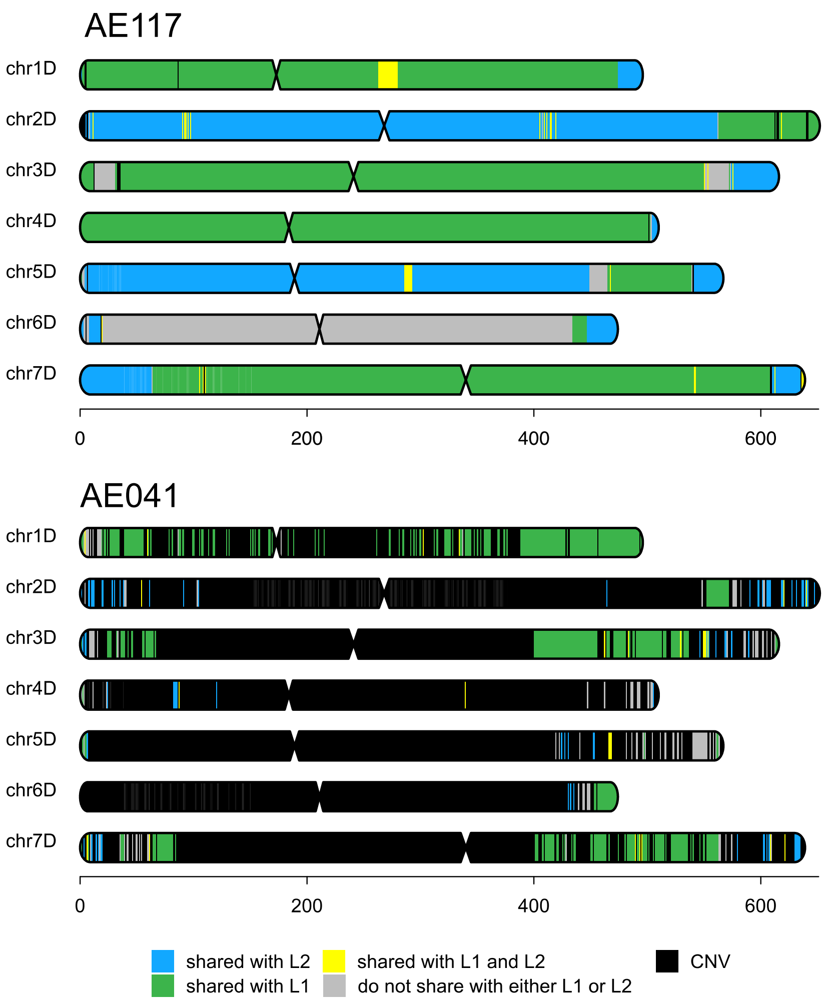


**Supplementary Fig. 9 Chromosomal distribution of ancestry for two putative *Ae. tauschii* L3 accessions.** Top: AE117. Bottom: AE041. Both accessions are actual hybrids between *Ae. tauschii* accession from L1 and L2.

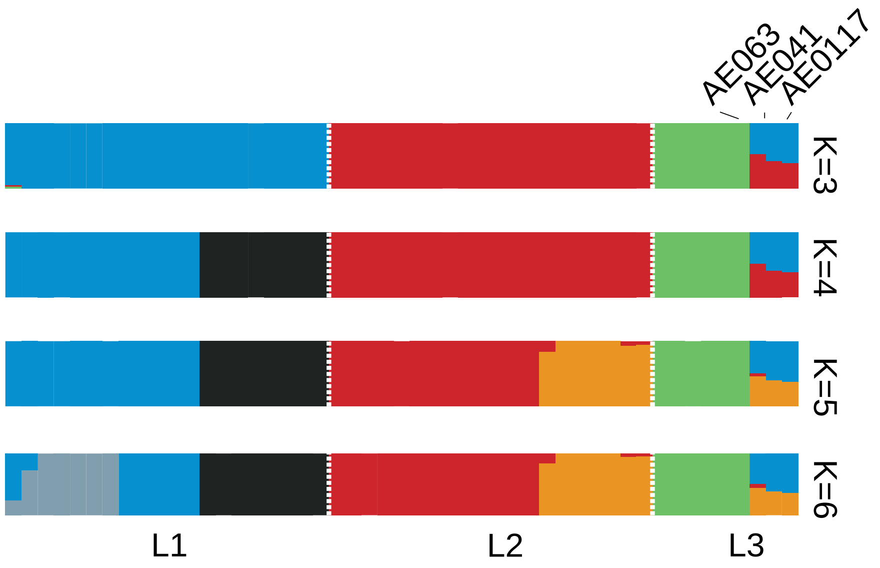


**Supplementary Fig. 10 Model-based clustering of the accessions from all *Ae. tauschii* three lineages.** 20 accessions were randomly selected from each of *Ae. tauschii* L1 and L2 for even sampling, along with the nine accessions of *Ae. tauschii* L3.

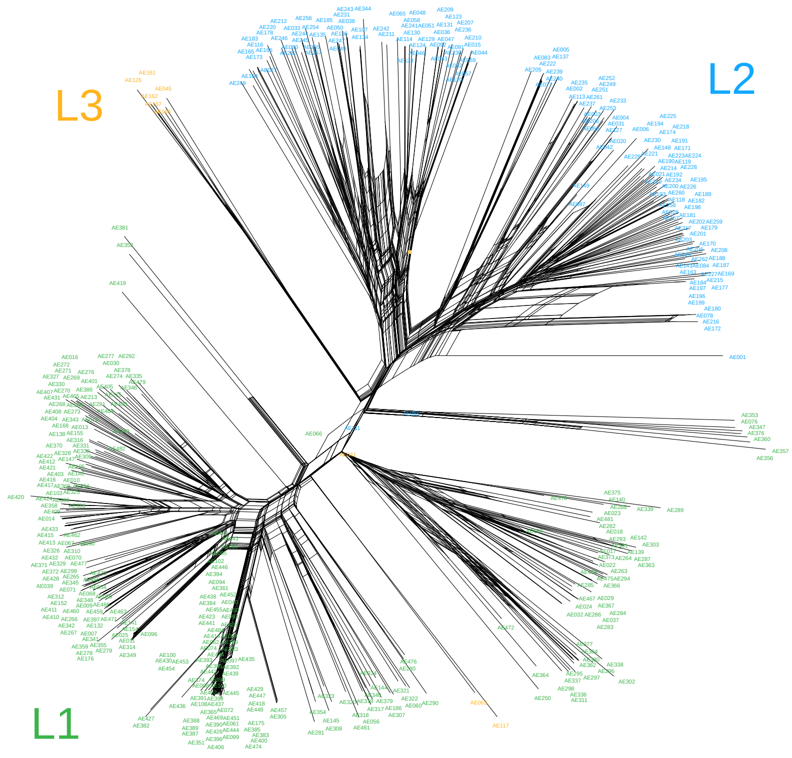


**Supplementary Fig. 11 Phylogenetic network of *Ae. tauschii* accessions using D genome AHG-based distance.** The network was constructed using NeighborNet (NNet) from SplitsTree4 (Huson and Bryant, 2005) based on D genome AHG distance. The reticulated events in the networks are represented by parallel branches.

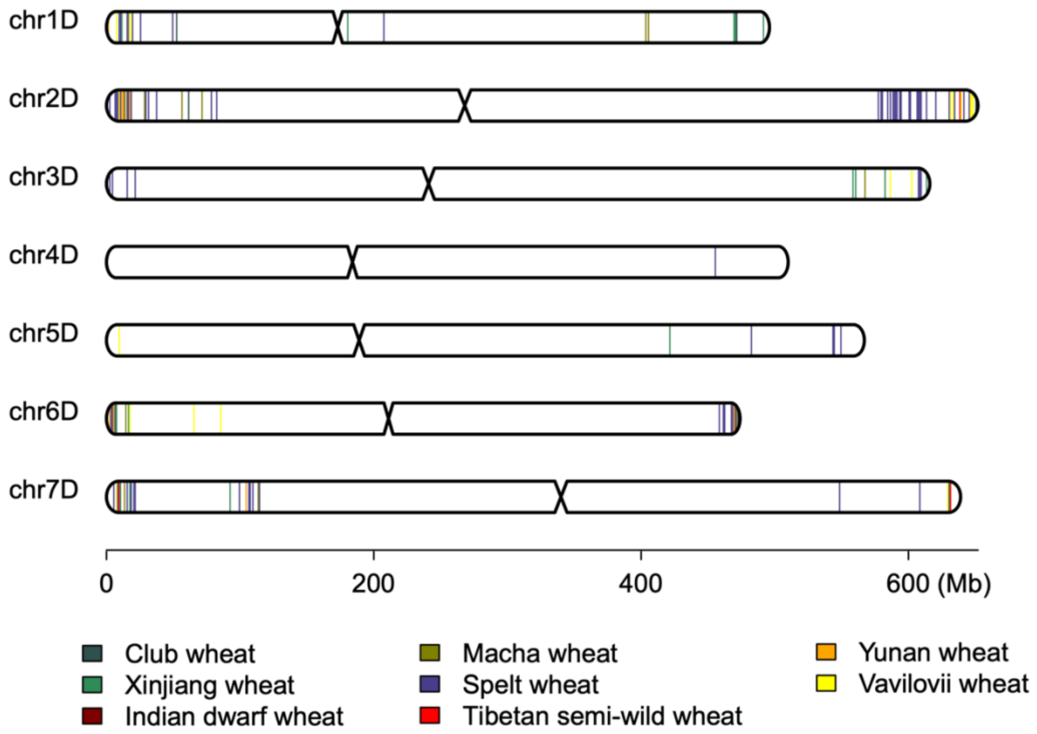


**Supplementary Fig. 12 Chromosomal distribution of AHGs specific to (sub)species of hexaploid wheat across D chromosomes.**

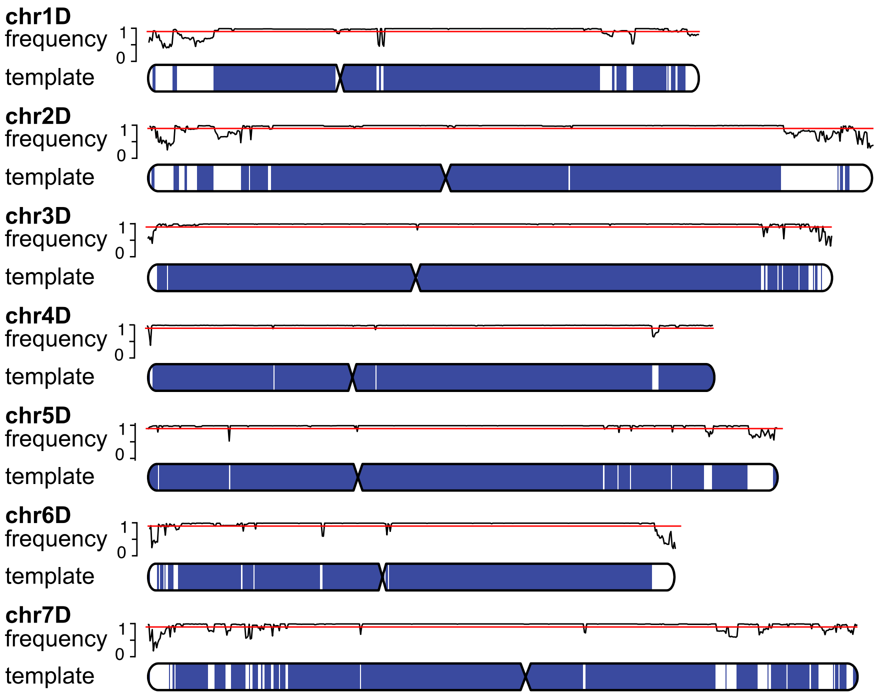


**Supplementary Fig. 13 Partition of the genetic diversity template region of hexaploid wheat D genome.** The frequency of dominant AHG is shown above seven D chromosomes. Red line, frequency of 90%. Blue blocks: template region.

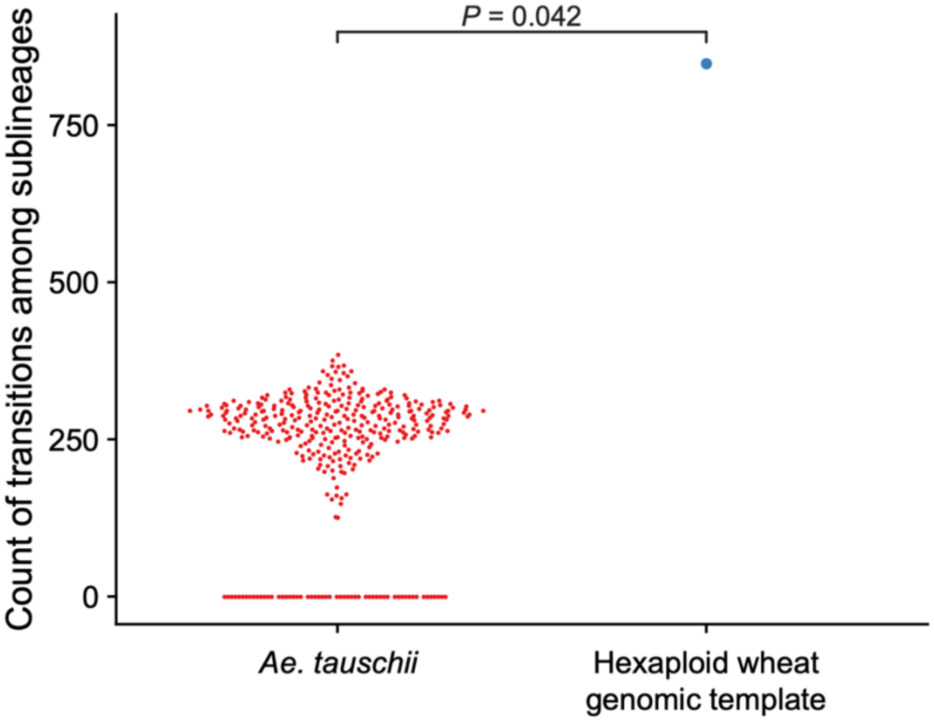


**Supplementary Fig. 14 Comparison of counts of ancestral transitions across D chromosomes between *Ae. tauschii* and the hexaploid wheat D genomic template accessions**. Transitions among sublineage origin between adjective genome windows were counted. Mann-Whitney U test was used.

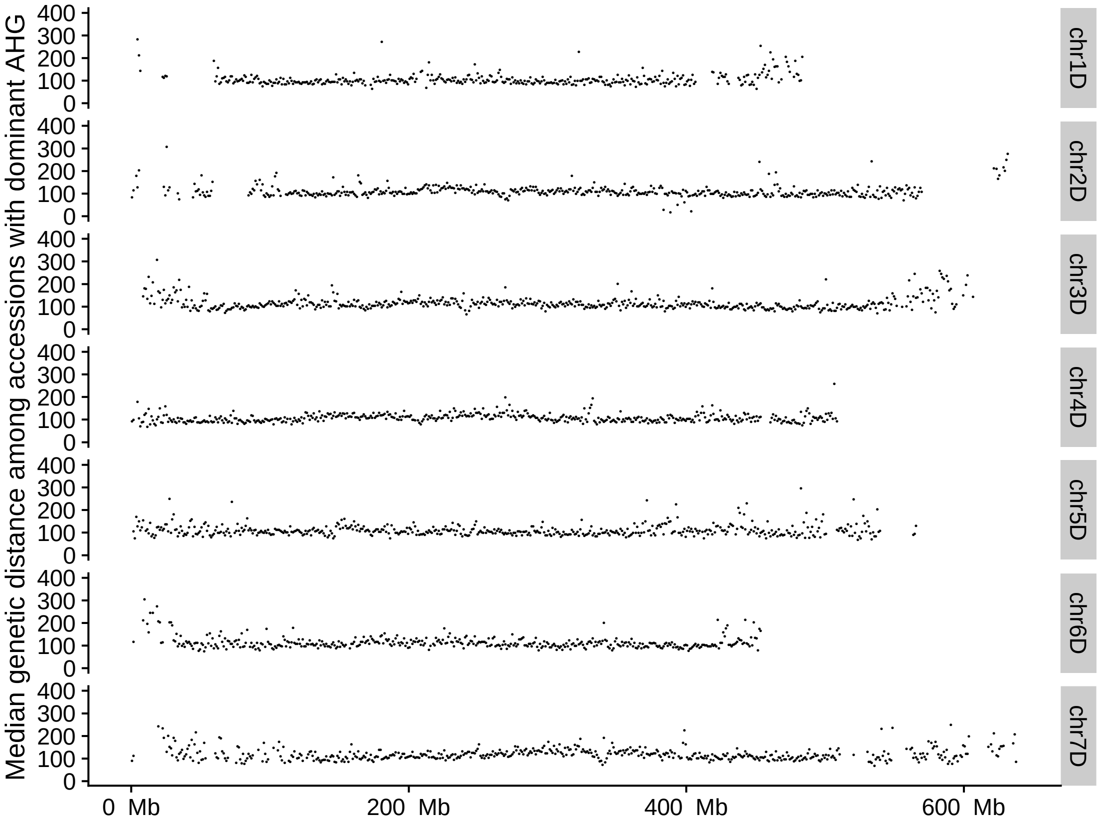


**Supplementary Fig. 15 Chromosomal distribution of genetic diversification levels among hexaploid wheat accessions across D chromosomes.** Genetic distance among accessions with dominant AHG was summarized to the median value for each 1-Mb genomic window.

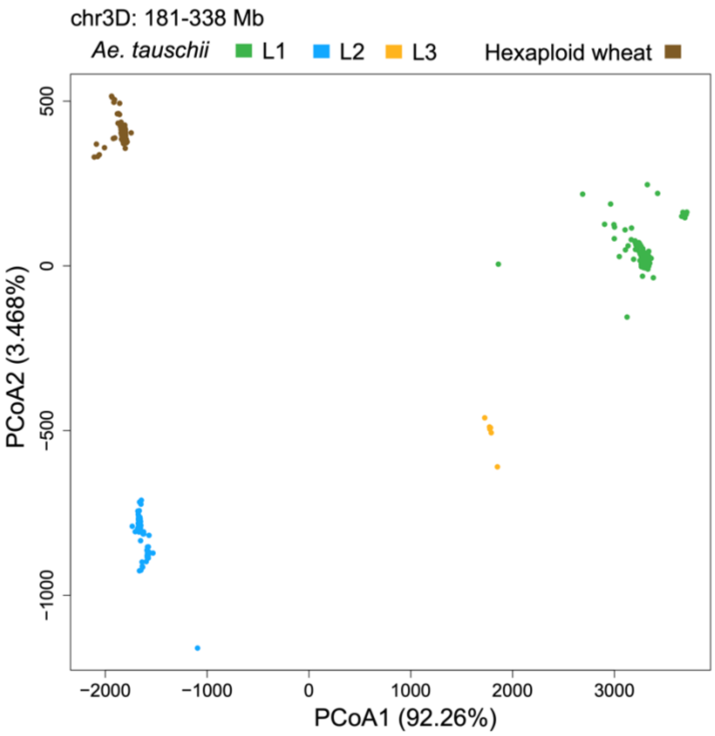


**Supplementary Fig. 16 Principal component analysis of *Ae. tauschii* and hexaploid wheat accessions for genomic region chr3D:181-338 Mb.** AHG-based distances for all 1-Mb windows in D genomes were used.

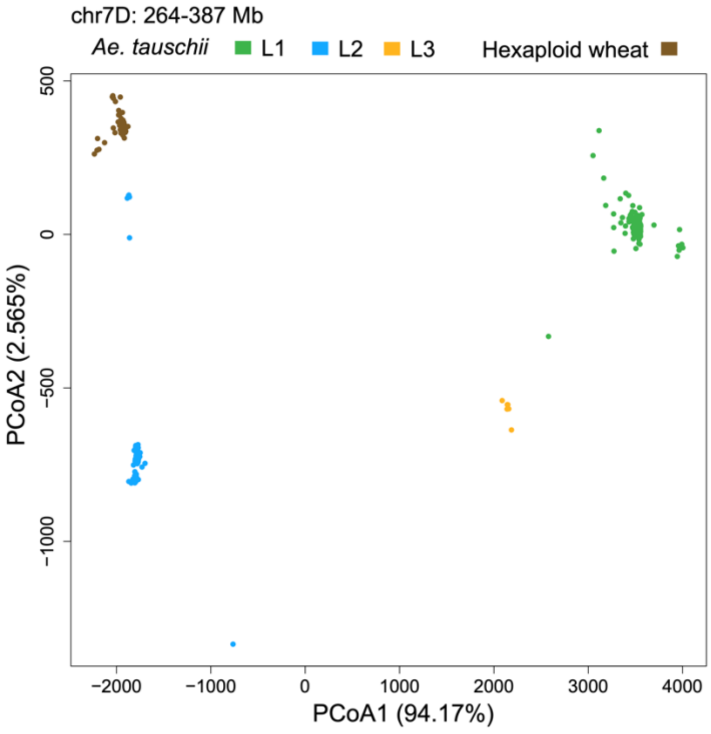


**Supplementary Fig. 17 Principal component analysis of *Ae. tauschii* and hexaploid wheat accessions for genomic region chr7D:264-387 Mb.** AHG-based distances for all 1-Mb windows in D genomes were used.

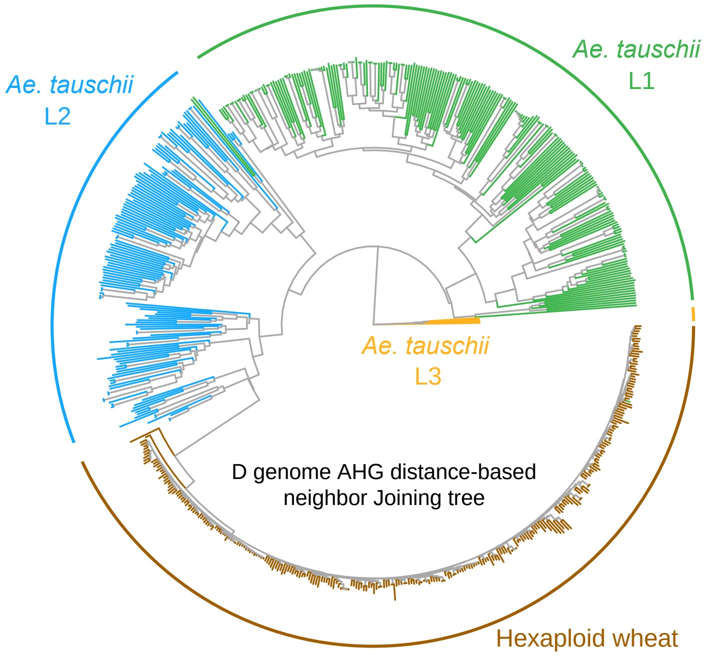


**Supplementary Fig. 18 D genome AHG distance-based neighbour-joining tree of *Ae. tauschii* and hexaploid wheat.** Four major taxonomic groups are marked by coloured lines along the circumference.

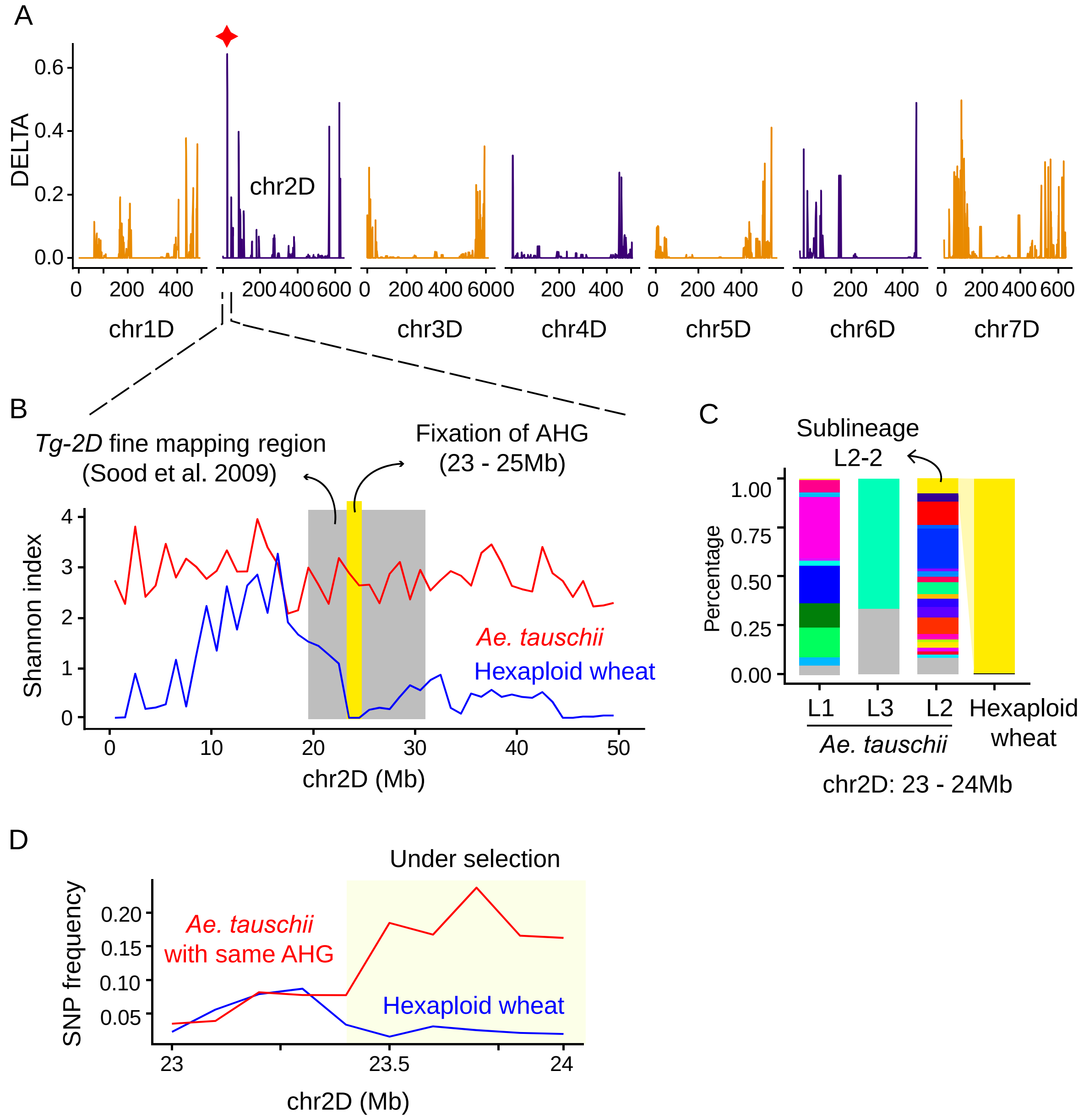


**Supplementary Fig. 19 Genomic diversity comparison and ancestry tracing of *Tg-2D***. A, The comparison of Shannon diversity index of AHG among adjective window as shown in DELTA statistics. The position of Tg-2D was labeled. B, The Shannon diversity index of AHG types of accessions from *Ae. tauschii* and hexaploid wheat across the first 50Mb of the 2D chromosome. The genomic region of *Tg-2D* fine mapping (Sood et al., 2009) was denoted with grey shading. Within the fine mapping region, the AHG type had been fixed across hexaploid wheat in chr2D:23-25Mb (yellow). C, Dynamics of AHG frequencies for chr2D:23-25 Mb block. The selected AHGs were connected through taxonomic groups with colored ribbons. The *Ae. tauschii* sublineage which accessions carrying selected AHG types were labeled with IDs. D, The binwise SNP frequency comparison between hexaploid wheat and *Ae. tauschii* with AHG type fixed in hexaploid wheat across chr2D:23-24 Mb. The frequency differs in region chr2D:23.4 - 24 Mb.

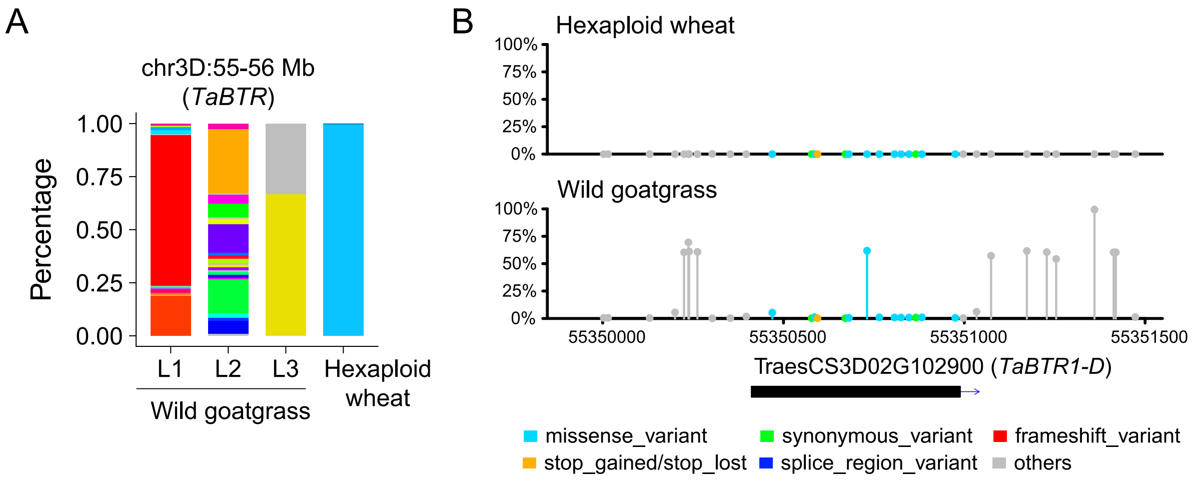


**Supplementary Fig. 20 Genomic diversity comparison and mutation detection of *TaBTR1-D*.** A, Dynamics of AHG frequencies for chr3D:55-56 Mb block. B. Lollipop graph with y-axis indicates mutation frequency in hexaploid wheat and wild goatgrass wheat accessions; mutation types were distinguished by different colors.

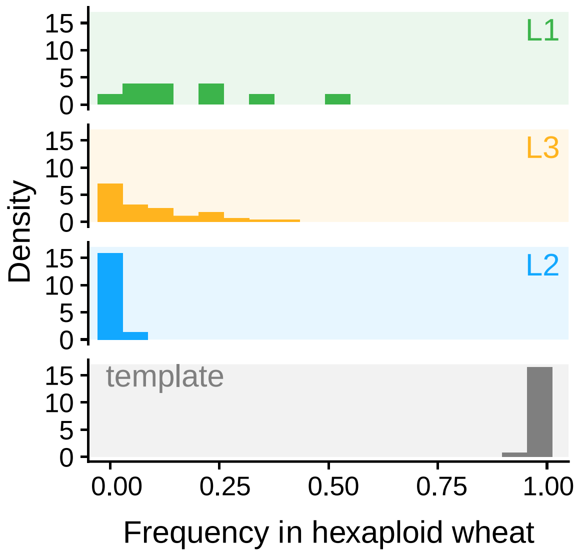


**Supplementary Fig. 21 Frequency distribution of non-template AHGs that could be traced back to L1, L3 and L3 in hexaploid wheat**. The frequency distribution of template AHGs is shown for comparison.

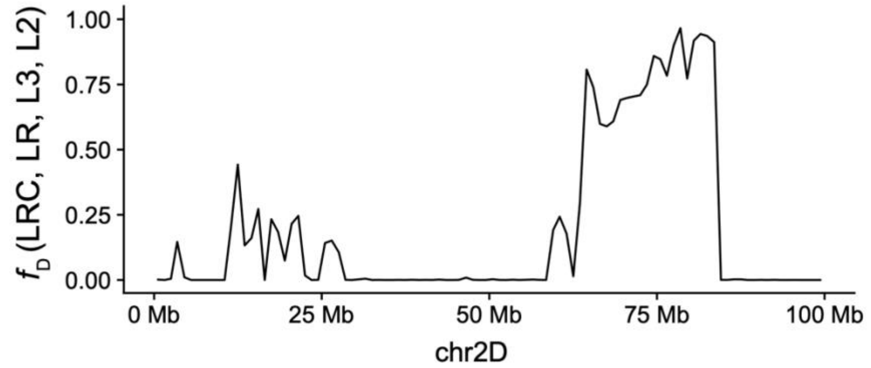


| Accession | Group | Accession | Group |
|---|---|---|---|
| GH397 | LR | GH474 | LRC |
| GH343 | LR | GH309 | LRC |
| GH284 | LR | GH217 | LRC |
| GH399 | LR | GH275 | LRC |
| GH405 | LR | GH161 | LRC |
| AE180 | L2 | AE162 | L3 |
| AE195 | L2 | AE157 | L3 |
| AE251 | L2 | AE158 | L3 |
| AE192 | L2 | AE126 | L3 |
| AE223 | L2 | AE160 | L3 |

**Supplementary Fig. 22 The distribution of $f_D$ statistics along the first 100Mb of 2D chromosome.** LRC, LR, L3, L2 were used as the P1, P2, P3, P4 of $f_D$ statistics, respectively. L2, *Ae. tauschii* subsp. *strangulata* (L2). L3, *Ae. tauschii* L3. LR, hexaploid wheat landraces of L3 introgression detected based AHG. LRC, hexaploid wheat landraces with no introgression detected. The containing accessions of each group are shown in the right table.

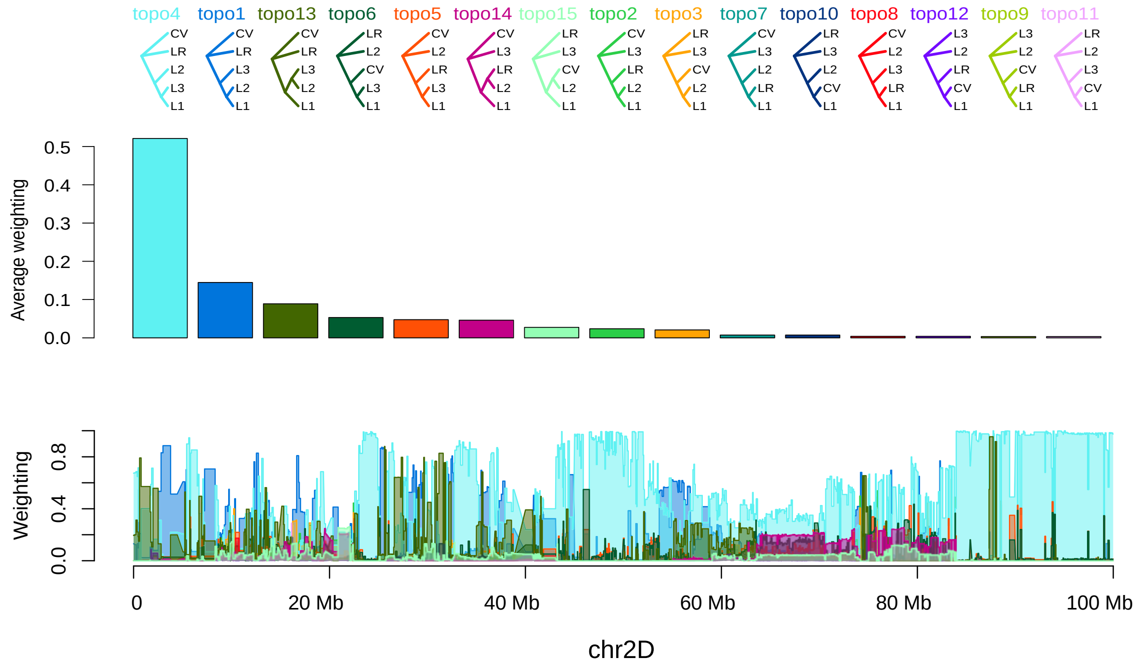


**Supplementary Fig. 23 Topology weighting statics across the first 100 Mb of chr2D.** The statistics were estimated using Twiss (Martin and Van Belleghem, 2017). The percentage of all subtrees matching each topology is shown by vertical bars.

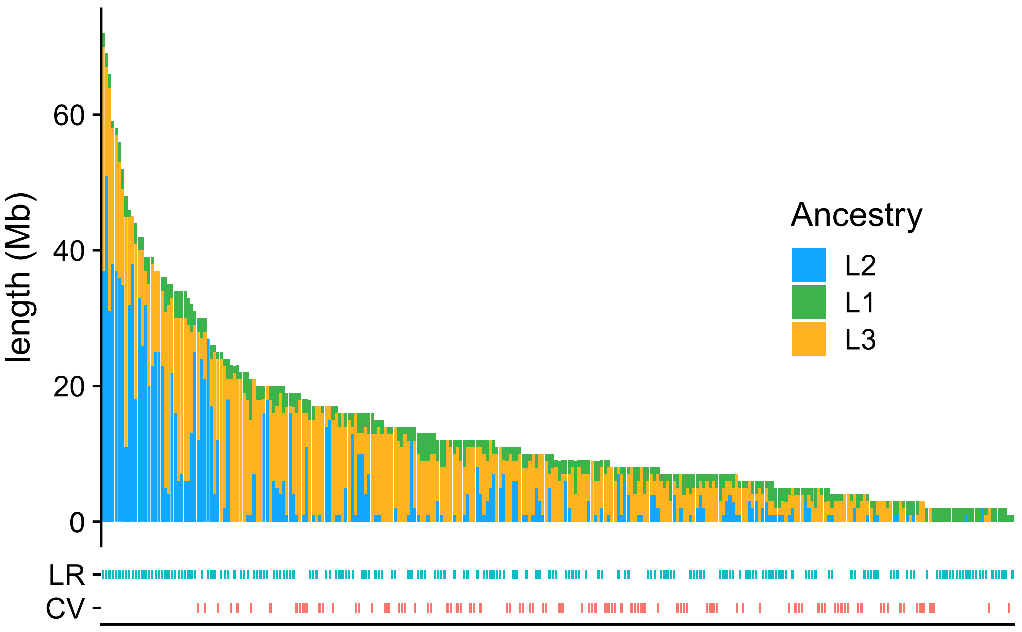


**Supplementary Fig. 24 Cumulative length of non-template AHGs that could be traced back to *Ae. tauschii* L1, L2 and L3 from hexaploid wheat.** The taxonomic group of each accession are shown.

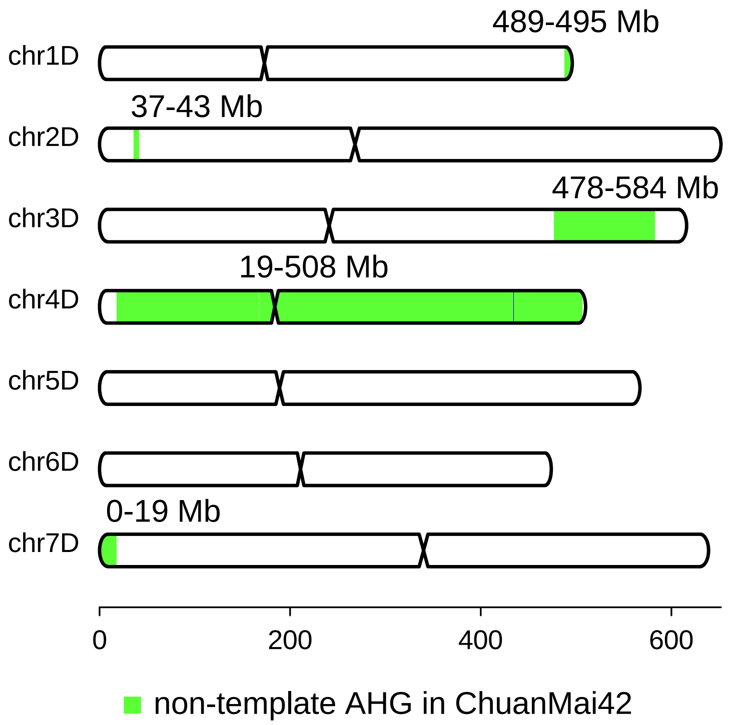


**Supplementary Fig. 25 Chromosomal distribution of non-template AHGs in hexaploid wheat cultivar CM42.** Five major genomic regions are annotated with green blocks and corresponding start and end positions.

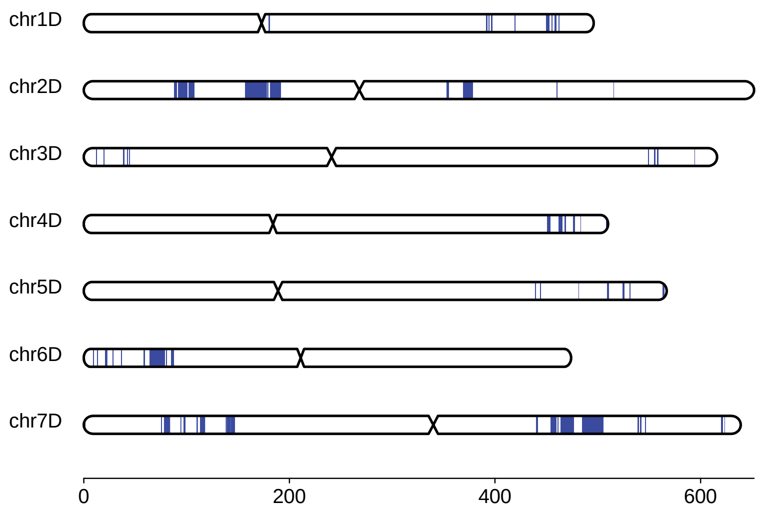


**Supplementary Fig. 26 Distribution of AHGs that could be traced back to L2 from hexaploid wheat.**

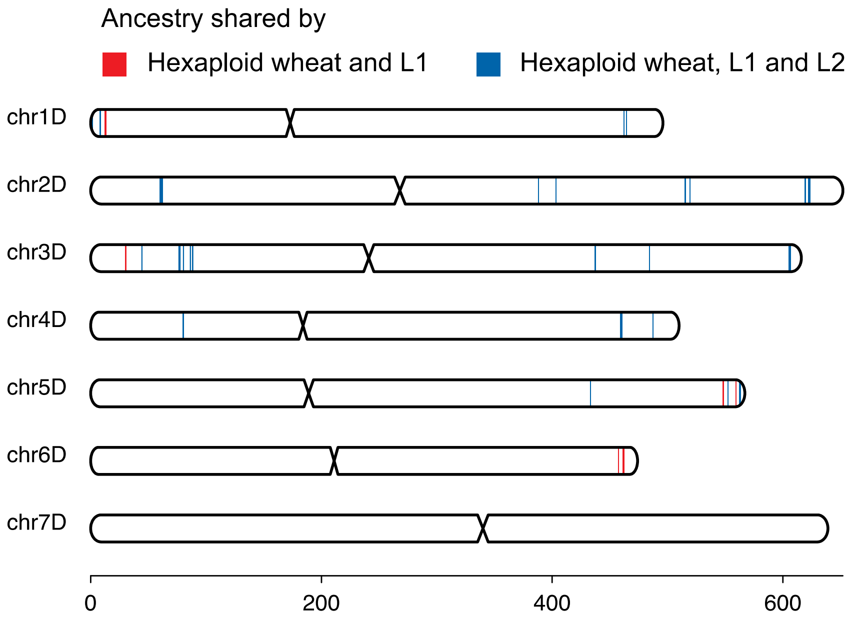


**Supplementary Fig. 27 Distribution of AHGs that could be traced back to L1 in hexaploid wheat.** AHGs that could be only traced back to *Ae. tauschii* L1 and AHGs that could be traced back to both *Ae. tauschii* L1 and L2 are labelled with red and blue colors, respectively.

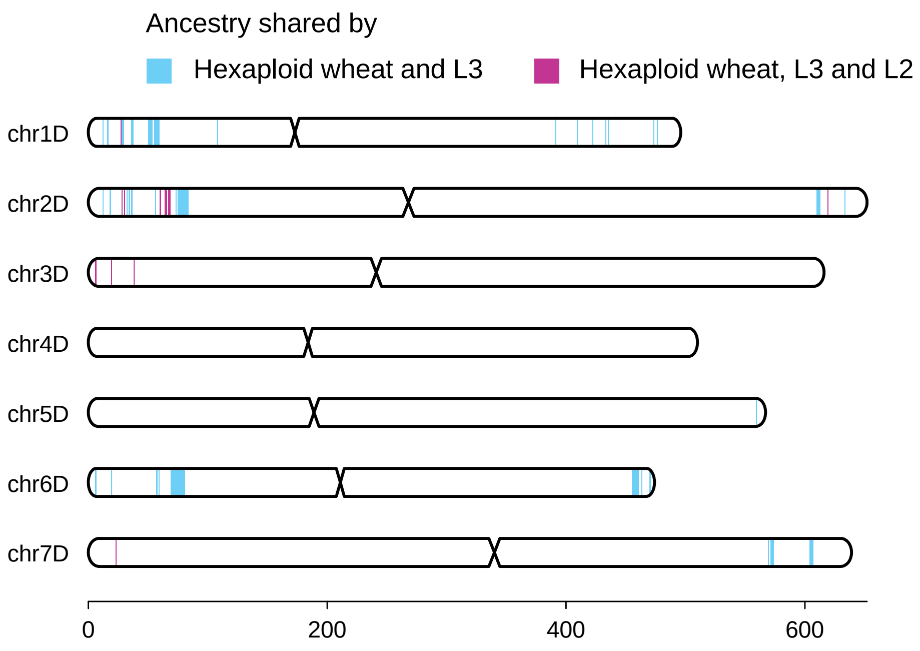


**Supplementary Fig. 28 Distribution of AHGs that could be traced back to L3 from hexaploid wheat.** AHGs that could be only traced back to *Ae. tauschii* L2 and AHGs that could be traced back to both *Ae. tauschii* L2 and L3 are labeled with blue and purple colors, respectively.

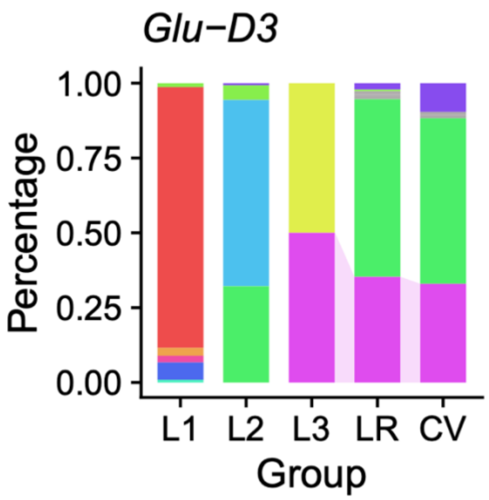


**Supplementary Fig. 29 Dynamics of AHG frequencies for the 1-Mb genomic window residing *Glu-D3*.** The selected AHGs were connected through taxonomic groups with colored ribbons.

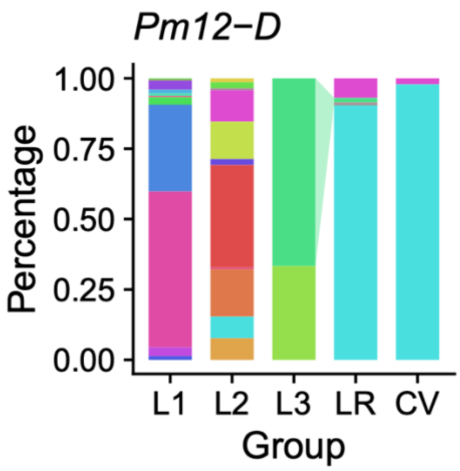


**Supplementary Fig. 30 Dynamics of AHG frequencies for the 1-Mb genomic window residing *Pm12-D*.** The selected AHGs were connected through taxonomic groups with colored ribbons.

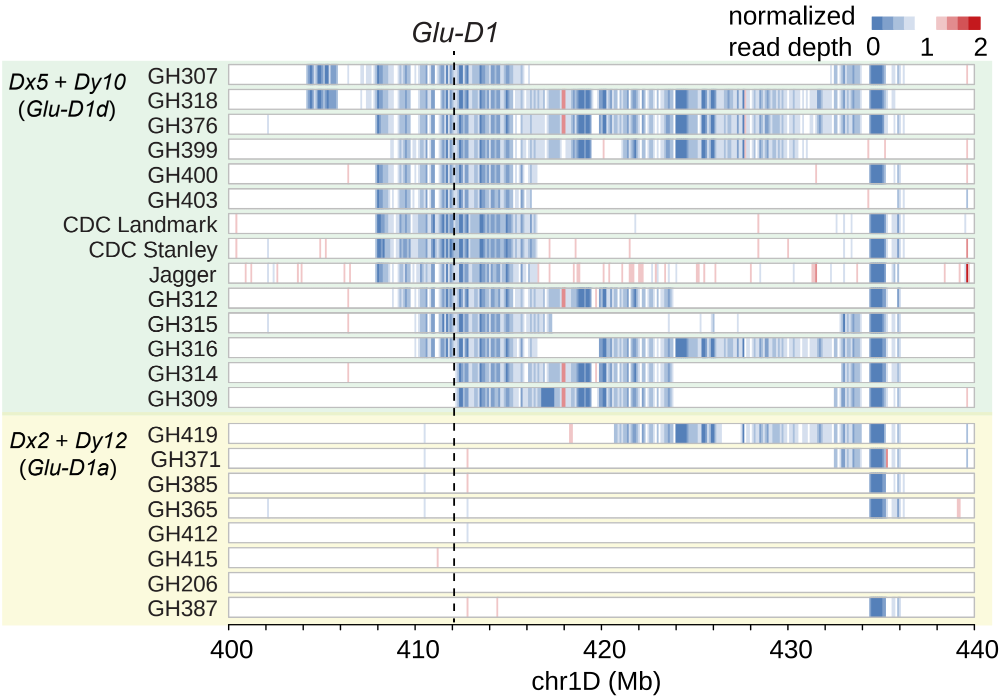


**Supplementary Fig. 31 Heatmap of normalized read depth in the flanking region of *Glu-D1* loci**. Genomic region chr1D: 400-440 Mb for 15 and 10 hexaploid wheat accessions with *Glu-D1a* and *Glu-D1d* genotypes are shown, respectively. The genomic position of *Glu-D1* loci is labeled.

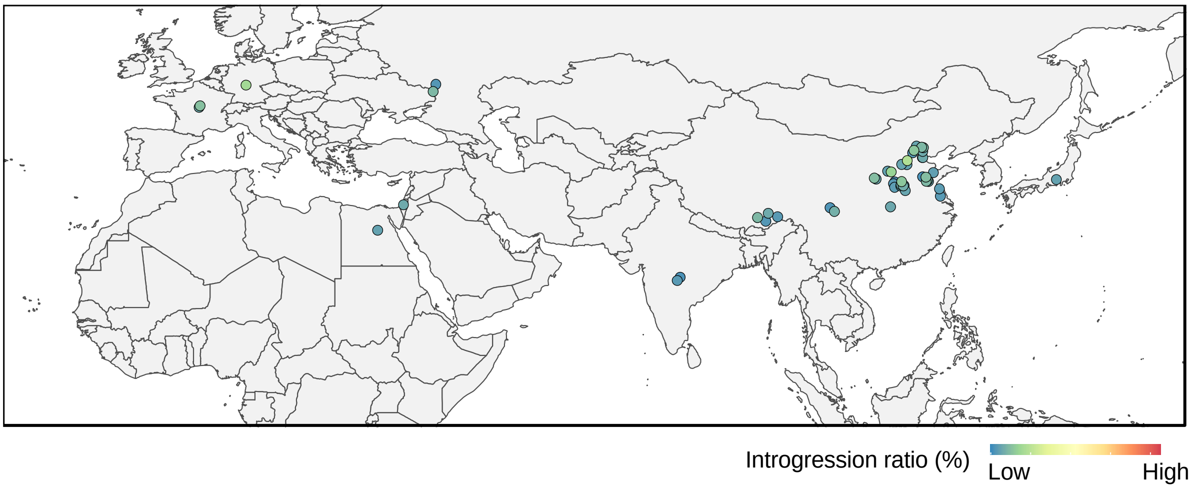


**Supplementary Fig. 32 Geographical distribution of hexaploid wheat cultivars with various non-template AHG ratios.** Dots represent individual bread wheat cultivar accessions.

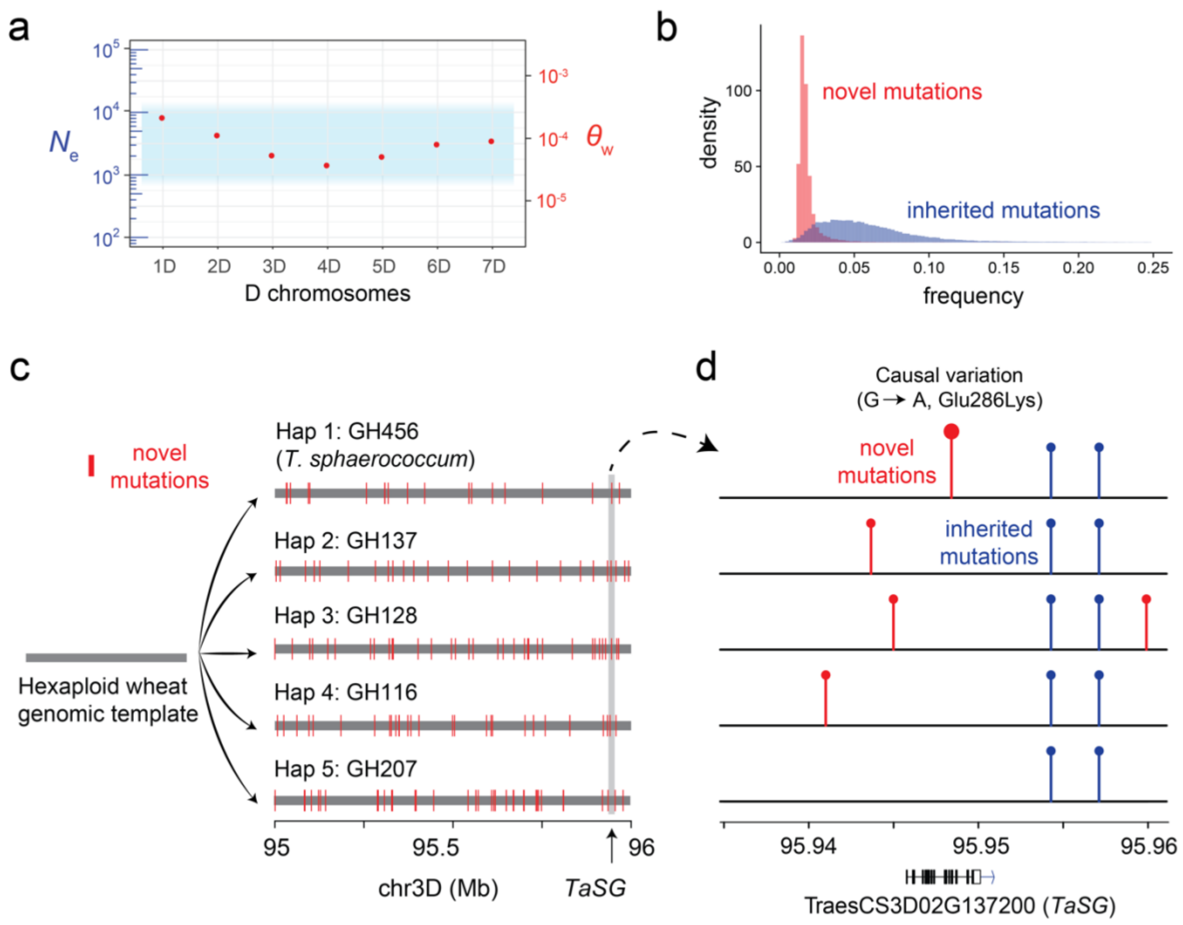


**Supplementary Fig. 33 Frequency and distribution of novel mutations in hexaploid wheat D subgenome.** a, Population diversity $\theta_w$ of novel mutations identified in this work is consistent with a population with an effective population size of the hexaploid wheat D genome (Zhao *et al.*, 2023). b, The frequency of novel mutations and inherited mutations. c, Five haplotypes of the genomic region (chr3D: 95-96 Mb) residing *TaSG* derived from the genomic template of hexaploid wheat D subgenome, with novel mutations labeled. d, Novel mutations and inherited mutations of five haplotypes aligned with the gene model of *TaSG*.

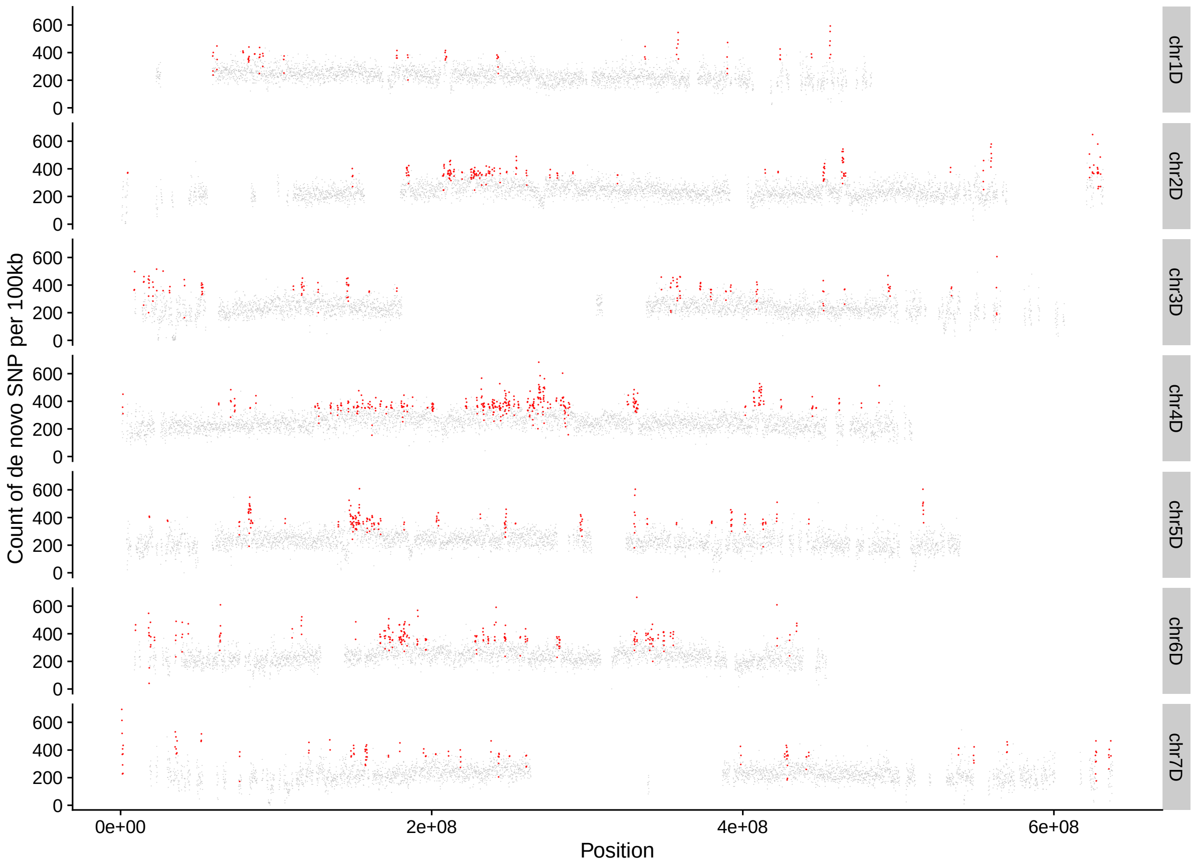


**Supplementary Fig. 34 Chromosomal distributions of the density of novel SNPs in hexaploid wheat D genome.** Hotspot regions were highlighted with red points.

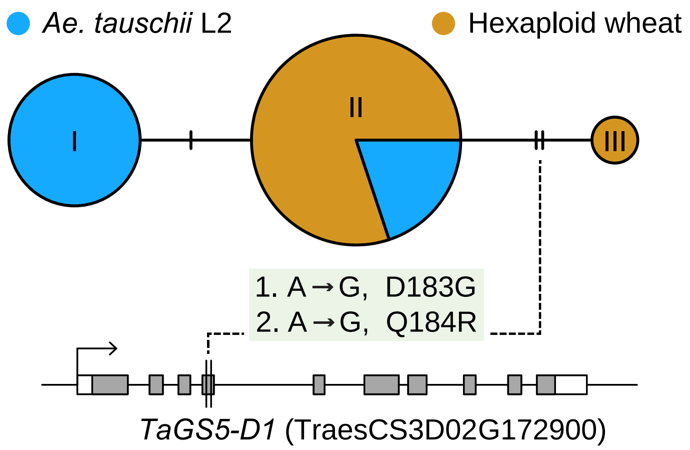


**Supplementary Fig. 35 The haplotype network and gene structure for the *TaGS5-D1*.** The gene structure highlights the location of two vital variants found only in hexaploid wheat, which result in the changes of the amino acid. The haplotypes were consecutively labeled using Latin numbers according to the evolutionary relationship.

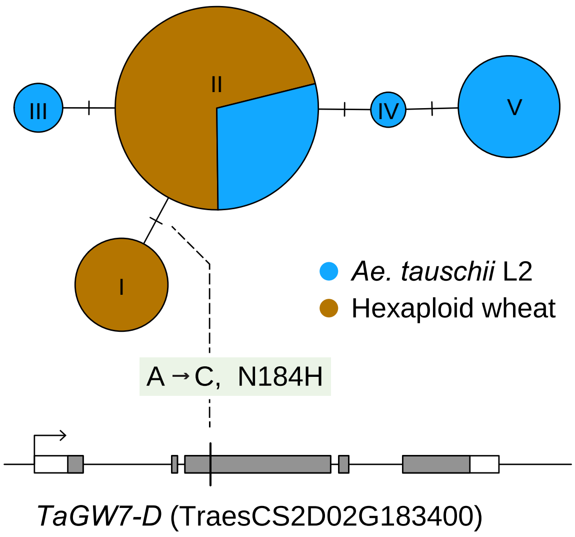


**Supplementary Fig. 36 The haplotype network and gene structure for the *TaGW7-D*.** The gene structure highlights the location of a vital variant found only in hexaploid wheat, which results in amino acid changes. The haplotypes were consecutively labeled using Latin numbers according to the evolutionary relationship.

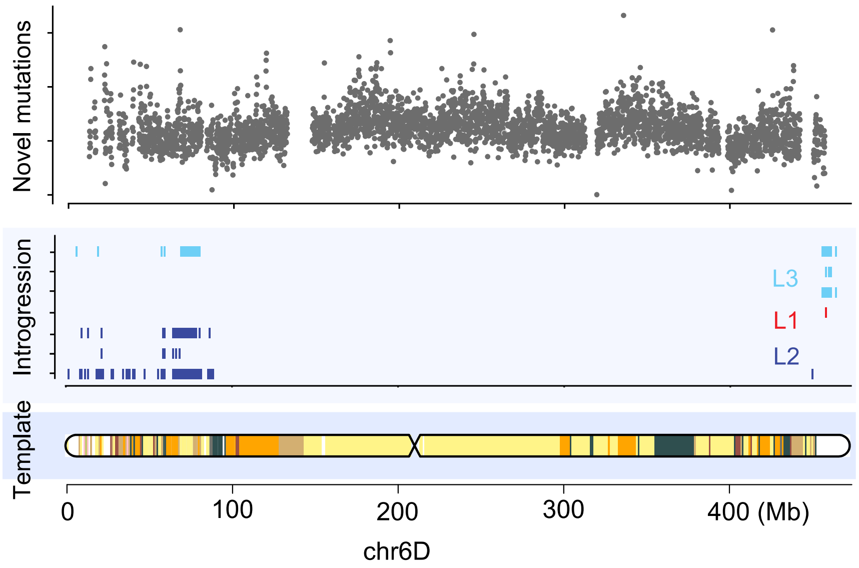


**Supplementary Fig. 37 Schematic diagram showing the inferred formation process of genetic diversity in hexaploid wheat D genome.** Hexaploid wheat shared the same set of genetic diversity template inherited from primary hexaploid wheat. Diversified introgressions from *Ae. tauschii* shaped the D genome after the hexaploidization event. Novel mutations further accumulated on the D genome along with spreading.

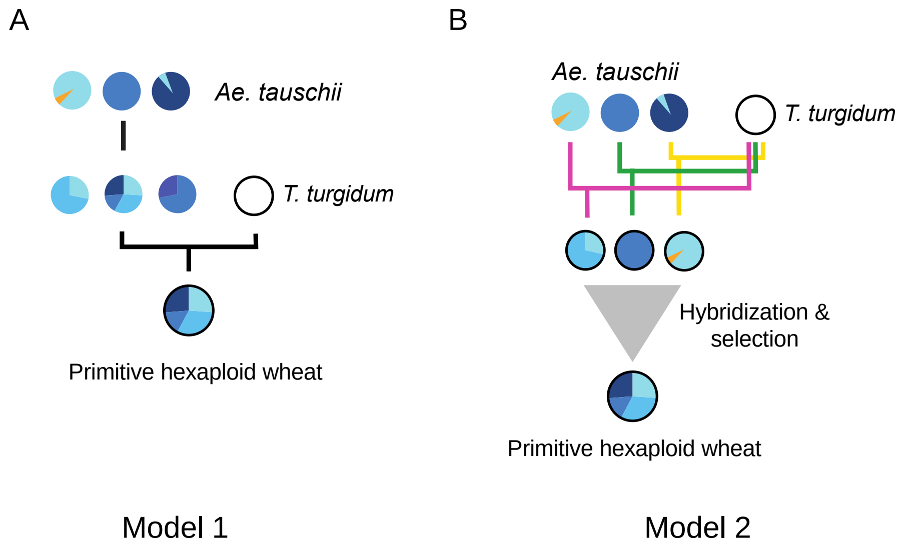


**Supplementary Fig. 38 The two putative evolutionary models leading to the ancestral mosaic genomic template of hexaploid wheat D genome.** A, Hexaploidization happened between tetraploid wheat and a high admixture *Ae. tauschii* accession. B, Hexaploidization happened multiple times between tetraploid wheat and various *Ae. tauschii* accessions. Hybridization happened among newly formed hexaploid wheat. Upon selection, only one hexaploid wheat thrived and replaced all other transitive hexaploid wheat.
